# Sex-allocation trade-offs and their genetic architecture revealed by experimental evolution

**DOI:** 10.1101/2024.12.11.627930

**Authors:** Jörn F. Gerchen, Nora Villamil-Buenrostro, Xinji Li, Ehouarn Le Faou, John R. Pannell

**Affiliations:** Department of Ecology and Evolution, University of Lausanne, 1015 Lausanne, Switzerland; Current address: Department of Botany, Faculty of Science, Charles University, 128 01 Prague, Czech Republic; Current address: Lothian Analytical Services, NHS Lothian, WaverleyGate 2-4 Waterloo Place, Edinburgh, EH1 3EG, Scotland

**Keywords:** life history, sexual system, sex ratio, hermaphroditism, dioecy, monoecy, leaky dioecy, quantitative trait locus

## Abstract

The theory of sex-allocation is hailed as one of evolutionary biology’s great successes. But while it has successfully predicted strategies of resource allocation to sons versus daughters under a wide range of conditions in species with separate sexes, it has been difficult to apply to hermaphroditic species because of the repeated failure to verify the fundamental assumption of a direct genetic trade-off between male and female allocations made by hermaphrodites. This assumption supposes that mutations that increase an individual’s male allocation reduce its female allocation in equal measure, yet there is almost no evidence for such ‘trade-off’ loci in hermaphroditic populations. Here, we use experimental evolution to generate wide variation in hermaphroditic sex allocation in an annual plant and demonstrate a clear trade-off between the two sexual functions within populations as well as over time, with populations evolving both increased male and reduced female allocation. Quantitative trait locus (QTL) analysis performed on multiple crosses further reveals the segregation of allelic variation at trade-off loci. Taken together, our results demonstrate a phenotypic trade-off in sex allocation in hermaphrodites and provide compelling evidence for sex-allocation trade-off loci, verifying the fundamental assumption of sex-allocation theory and securing its general applicability to both dioecious and hermaphroditic species.

## Introduction

Sex-allocation theory, which predicts strategies of resource allocation to sons versus daughters, is hailed as one of evolutionary biology’s great successes (West et al. 2000, Frank 2002). Prime reasons for its empirical success include the relative ease with which the sex ratio can be measured in species with separate sexes, and the fact that the theory’s fundamental assumption concerns a direct allocation trade-off between the production of sons versus daughters (Fisher 1930, Charnov 1982, Frank 1986, Hardy 2002, West 2009). Sex-allocation theory is also very general, in the sense that its predictions can easily be extended to hermaphroditism in terms of investment into male and female functions (e.g., the production of eggs versus sperm, ovules versus pollen, male flowers versus female flowers, etc.), rather than only in terms of investment into sons and daughters of dioecious species (Charnov et al. 1976, Charlesworth and Charlesworth 1981, Charnov 1982, Goldman and Willson 1986, Charlesworth and Morgan 1991, Morgan 1992, Zhang and Jiang 2002). However, the theory’s predictions have been much less successfully tested in hermaphroditic populations than in those that are dioecious (Goldman and Willson 1986, Campbell 2000, Ashman and Majetic 2006, Schärer 2009), in large part because the fundamental trade-off assumption is so difficult to verify for hermaphrodites.

Numerous studies conducted on both plant and animal systems have sought evidence for an allocation trade-off between male and female functions, whether among individuals or at the modular level within individuals (reviewed in Ashman 2003, Schärer 2009). While some studies have documented elements of the expected trade-off (e.g., Raimondi and Martin 1991, Devisser et al. 1994, Trouvé et al. 1999, Koelewijn 2003), others have reported no trade-off (e.g., Petersen 1990, Eckhart 1993, Hoch and Levinton 2012, Varga and Kytöviita 2017) or even a positive rather than a negative correlation between male and female allocation (Campbell 2000, Locher and Baur 2000, Ashman 2003, Scharer and Ladurner 2003, Schärer 2009). A sex-allocation trade-off may simply not exist for some hermaphrodites if their male and female functions draw on different resources (Charnov et al. 1976, Cruden and Lyon 1985, Goldman and Willson 1986, Ashman 1994). For instance, pollen production may be limited by the availability of nitrogen and ovules and seeds by carbon (e.g., Harris and Pannell 2008), or male and female allocations may be made at different times (Bawa 1982, Herrera 1982, Goldman and Willson 1986, Schärer et al. 2005).

Alternatively, a genuine sex-allocation trade-off may be obscured by variation among individuals in other traits (Roff and Fairbairn 2007), e.g., with larger and thus better-provisioned individuals allocating more to both their male and female functions (van Noordwijk and de Jong 1986, Petersen and Fischer 1996, Fenster and Carr 1997, Trouvé et al. 1999, Campbell 2000, Schärer et al. 2001, Schärer et al. 2005). Attempts to expose trade-offs by limiting variation resource availability among individuals by feeding them a low-nutrient diet have also been met little success (Schärer et al. 2005).

Testing the assumption of a trade-off between male and female function requires a population with sufficient variation in sex allocation among individuals, yet we should expect selection often to have optimised sex allocation and thus to have removed variation among individuals (Fisher 1930, Barton and Turelli 1989). In principle, tests of the trade-off assumption could be performed by experimentally manipulating allocation, e.g., by preventing allocation through the removal of anthers or carpel buds on flowers before at the bud stage (Harris and Pannell 2008), or by preventing allocation to male function by severing the nerve to the male gonads of hermaphroditic snails (Devisser et al. 1994). Such experiments have revealed modest evidence for trade-offs, but the manipulations can also have confounding physiological effects that compromise interpretation of allocation measurements in terms of trade-offs (Schärer 2009). Overall, the limited success we have had in demonstrating sex-allocation trade-offs in hermaphrodites limits the extent to which we can use sex-allocation theory to predict reproductive strategies for a significant number of species on Earth (Schärer 2009).

Our failure to demonstrate sex-allocation trade-offs can also be attributed to our limited understanding of the genetic architecture of allocation. Sex-allocation theory implicitly assumes a genetic architecture in which mutations that increase allocation to one sexual function cause a decrease in allocation to the other (Charnov et al. 1976, Charnov 1982, West 2009), with alleles potentially segregating at what we might term ‘sex allocation loci’. Indeed, theoretical modelling of the evolution of sex allocation explicitly considers the fate of mutations at such loci (e.g., Charnov et al. 1976, Charlesworth and Charlesworth 1981, Lesaffre et al. 2024a, b). Yet while the quantitative genetic architecture of sex-ratio variation has been investigated for several dioecious species of vertebrates and insects (reviewed in Pannebakker et al. 2020), we remain completely ignorant of the genetic architecture of sex allocation in hermaphrodites. Ramm et al. (2019) discovered genes in a flatworm whose gonad-specific expression was associated with either increased testis size or reduced ovary size under social conditions predicted to favour a shift to greater male allocation, but the genes appear to act independently, and the study’s results do not imply the existence of any direct allocation trade-off. We thus continue to lack evidence for both a phenotypic trade-off in sex allocation and the assumed antagonistic pleiotropic effects of mutations on both male and female allocations, as assumed by theory.

Here, we demonstrate a strong sex-allocation trade-off in an annual plant with both male and female flowers as well as evidence for the segregation of alleles at ‘sex-allocation loci’ with antagonistic pleiotropic effects. The case is particularly noteworthy, as the male and female functions of the species draw on partially different resources, such that expectations for a direct trade-off are reduced: pollen production draws heavily on nutrients such as nitrogen, while seed and fruit production are likely limited by carbon (Harris and Pannell 2008). We demonstrate both a negative correlation between male and female allocations among individuals within populations as well as through divergence in sex-allocation in populations evolving under experimental evolution over time. We further characterise the genetic architecture of the sex-allocation trade-off from a quantitative trait locus (QTL) analysis based on crosses between extreme genotypes from our experiment; these crosses reveal two classes of genomic regions associated with sex allocation: those segregating for variation in male or female allocation independently of one another (and thus inconsistent with trade-off assumptions); and those segregating at trade-off loci for ‘plus’ and ‘minus’ alleles with opposite effects on male and female allocations. The loci that explain the greatest amount of variation in sex-allocation are of the latter type and can be found on different chromosomes, suggesting that a diverse set of genetic variants with effects on sex expression can segregate in natural populations. Our results represent a significant advance for applying sex allocation theory to hermaphrodites, and they strengthen the empirical basis for quantitative models of evolutionary transitions between hermaphroditism and dioecy.

## Materials and Methods

### Overview of approach, and study populations

We studied variation in sex allocation and associated genetic variation in a subset of six populations of the wind-pollinated annual plant *Mercurialis annua* that have been evolving under semi-natural conditions in gardens in Lausanne, Switzerland, for 10 years under two treatments with contrasting sex allocation and mate availability (Cossard et al. 2021). Diploid populations of *M. annua* are dioecious, with an XY sex-determination system and a 1:1 sex ratio (Russell and Pannell 2015, Veltsos et al. 2018, Veltsos et al. 2019). Females produce their pistillate flowers and fruits on short pedicels in the leaf axils, while males produce staminate flowers on erect inflorescence stalks (peduncles). In our experiment, three populations (labelled 2, 4 and 6) comprised a 1:1 sex ratio of XY males and XX females, as in wild populations; these are labelled the ‘Control’ populations. Three populations (labelled 1, 3 and 5) were established from the same ancestral seed population as the Control populations, but comprised only XX females at the start of the experiment, following the removal of all males. Because the removal of males meant that these populations lacked a Y chromosome, they are labelled ‘X-only’ populations.

As is the case for many dioecious plants (Ehlers and Bataillon 2007), XX females of *M. annua* are occasionally inconstant or ‘leaky’ in their sex expression, producing a few fertile male flowers (Cossard and Pannell 2019). XX individuals in the X-only populations began the experiment as females, identical to those in the Control populations, but have evolved dramatically increased male flower production over the course of the experiment in response to strong selection on the sex allocation, as described by Cossard et al. (2021). In contrast, XX individuals co-occurring with males in the Control populations have maintained their strongly female sex allocation. The experiment has been propagated annually since 2012, with seeds sown in spring and the seeds they produce harvested in early summer and used both for resowing in the next generation and for downstream experiments and analysis. Further details of growth conditions in the experimental gardens are given in Cossard et al. (2021).

We generated in-depth phenotyping results of plants from one experimental X-only population from the latest generation as well as a multi-generational common garden based on stored seeds to demonstrate phenotypic evidence for trade-offs in sex allocation, both within and among X-only populations. We further explored the underlying genetics of these trade-offs using crosses between Control and X-only populations based on earlier generations of the experiment.

### Within-population variation in sex allocation

As a first test for a sex-allocation trade-off, we compared the male and female reproductive investment among individuals from one of the X-only populations (Population 5) at generation G11 (2022) of the experiment. The plants were harvested from the garden, placed in a fridge, and phenotyped within two days of harvest. We counted the number of fruits and the number of male flowers in the reproductive active apex (the top 20 cm) of each plant, then weighted the corresponding shoots and branches following drying at 40° C.

### Common garden experiment

To characterise the temporal dynamics of the trade-off in sex allocation between the female and male function from experimental all-female populations, we established a large common garden by growing seeds from all six populations retained from six generations (generation 1, 3, 5, 6, 7 and 8). All plants were grown together under same conditions and were exposed to the same pollination environment. On average, the common garden contained 31 plants per generation and per population (total *N* = 1116 plants). Plants were first established from seeds in seedling trays and then grown individual in pots with soil (140 Ricoter soil) with a slow-release fertiliser (Hauer-Tardit 3M: 500 g per 100 L of soil), arranged in a randomised block design in an open polytunnel, with each table treated as a representative block that contained replicate plants from all populations and all generations. Plant positions on each table were assigned to rows and columns using random numbers. Plants were grown over the spring and summer and harvested 20 weeks after germination. Plants suffered damage from thrips and viral infections and experienced a brief period of drought due to a failure in the automatic watering system over a weekend. Plants were allowed to recover fully before being measured, but some individuals were lost to disease and drought. Since phenotyping took over a month to complete, the last phenotyped plants had more time to grow than the first; we thus recorded the phenotyping date for each plant and included this as a co-variable in the analysis. We harvested plants from the common garden by cutting them at their base, just above the soil, and measured their height. We determined the sex allocation of each plant, focusing on a 30 cm sub-sample of each plant. On this sub-sample, we harvested all male flowers using tweezers, dried them in paper envelopes, and weighed them. The sub-sample was then kept in a porous plastic bag to dry out, promoting the release of seeds from fruits. The collected seeds were then weighed, along with the dry sub-sample, using a digital balance. The rest of the plant, dried in the same way, was also weighed. The plant’s ‘slenderness’ was calculated as the plant’s height divided by its biomass (Van Der Werf et al. 1995). Although plants from the Control populations were grown with plants from the X-only populations in this common garden, we focus here solely on the data collected from the X-only populations (1, 3 and 5).

### Statistical analyses of phenotypic variation in sex-allocation

With data from the generation G11 of Population 5, we tested whether investment in male flowers (pollen production) by females came at the cost of investment in seed production using a standardized major axis regression, such that any trade-off coefficient should be equivalent for both male or female trait as the response variable (REF). With data from the common garden, we performed a zero-inflated Gamma generalized linear model. This analysis included only females from the three X-only populations (with only XX individuals). The model therefore considers the effect of seed mass, generations, total height and slenderness (all fixed factors) on the mass of male flowers (response). Population, block and phenotyping date were included as random effects. Within-population variation in sex allocation was analysed similarly. Prior to performing the regression, both the number of fruits and the number of male flowers was standardized by dividing by the corresponding dry mass of the sub-sample to reduce the variance due to the quantity actually invested by the individuals in the sub-sample, and increase the power available to measure this trade-off (Van Noordwijk and De Jong, 1986); this standardized measure corresponds to an estimate of the ‘reproductive effort’ via female and male functions, respectively.

### QTL analysis

To map QTL for sex allocation in our experimental populations, we first generated four *F*_1_ crosses between plants from the X-only and Control populations from the fifth generation of the experiment. Three of these crosses were between individuals of the fifth generation of Control and X-only populations, using paternal parents from each of the three X-only populations. In addition, for a fourth cross we used a paternal plant from an additional individual from X-only Population 1 in which only 25% of the top pollen producers had been used to establish the generation prior to sampling, because this population showed particularly high male-flower production by the XX females. For each cross, we used one individual from a X-only as a paternal pollen donor and one female from a Control population as maternal ovule donor. We then grew the *F*_1_ plants from seeds and self-fertilized them to generate *F*_2_ plants. We finally phenotyped and genotyped the resulting *F*_2_ plants to estimate the biomass of male flowers (*W*_m_), female flowers and fruits (*W*_f_), and the whole plant (*W*_p_), and we defined as key variables of allocation to reproduction male reproductive effort (MRE) as *W*_m_/(*W*_m_ + *W*_f_ + *W*_p_)), female reproductive effort (FRE) as *W*_f_/(*W*_m_ + *W*_f_ + *W*_p_)), and relative male allocation (rMA) as *W*_m_/(*W*_m_ + *W*_f_).

We regressed out three variables of allocation on genotype data generated using a modified ddRad protocol (Peterson et al. 2012). We used the demultiplexed Illumina reads in an integrated analysis pipeline implemented in a reproducible Snakemake workflow ((Mölder et al. 2021), https://github.com/jgerchen/qtl_mapping), to build linkage maps and run QTL-analyses for each cross separately (see Supplementary Methods). In contrast to using a single analysis that incorporates all crosses, this approach allowed us to test specifically how a sex allocation trade-off is implemented in each paternal pollen donor for each cross and to compare the underlying architecture between experimental populations.

## Results

### Trade-off between male and female allocations and their evolution over time

Sex allocation in the common garden evolved towards greater male allocation over time, as already revealed by Cossard et al. (2021). This ongoing evolution of greater male allocation has come at cost of female allocation, with a clear trade-off between allocation to male and female functions in generation G11 of Population 5 (Figure 1). The slope coefficient relating allocation towards male function to that towards female allocation was -9.26 (95% CI [- 10.23; -8.38]), pointing to the equivalence of one female flower or fruit to about nine male flowers (Smith 2009, Friedman et al. 2013) (Figure 1). Data from the common garden confirmed the presence of a trade-off, the slope coefficient for this one being -1.54 (95%CI [-1.72; -1.38]), indicating a trade-off equivalence between one biomass unit of fruits and seed and 1.5 units of male flowers (Table 1). The common garden experiment also shows that mean allocation to male function per individual significantly increased over time (Figure 2), with each consecutive generation having associated negative effects on female flower biomass (Table 1). The sex-allocation trade-off affected taller and shorter plants similarly (Table 1). The significant negative effect of plant slenderness on male allocation indicates that the most elongated individuals tended to invest less in male flowers (Table 1).

**Figure 1.**
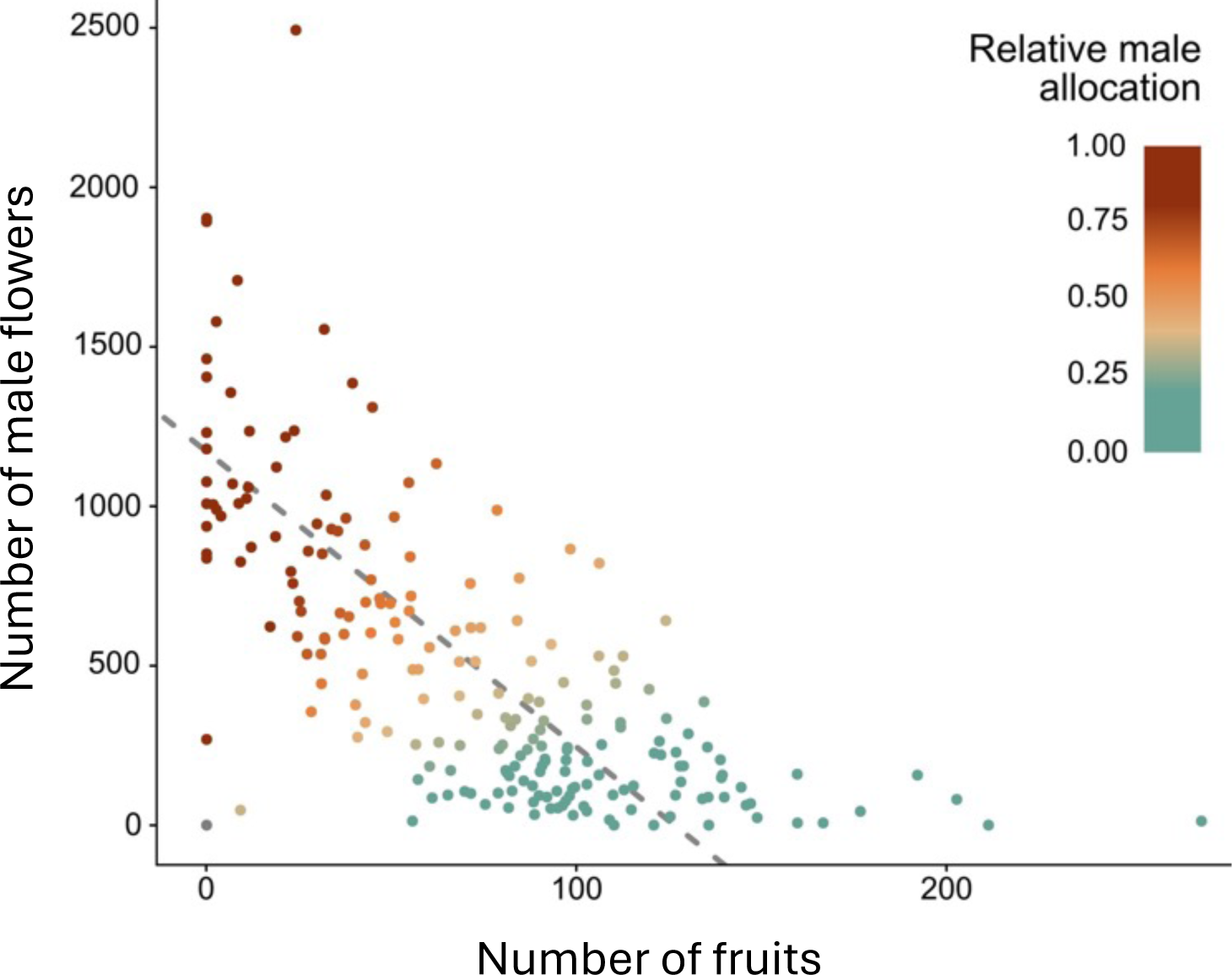
Relationship between the number of male flowers and the number of flowers and fruits counted on the sub-sample of one of the three X-only populations (Population 5, with only XX individuals). Both variables are standardised by the dry biomass of the sub-sample. The dotted line shows the result of the standardised major axis regression (intercept = 1171, 95%CI [1088; 1255], slope = -9.26, 95%CI [-10.23; -8.38], R2 = 0.47). The colour of the points reflects the relative male allocation in terms of numbers (analogous to rMA for biomass), computed as the standardized number of male flowers divided by the sum of the number of male flowers with that of female flowers and fruits. The calibrated female trait was calculated as the negative of the female trait divided by the slope (Klinkhamer et al. 1997).

**Figure 2.**
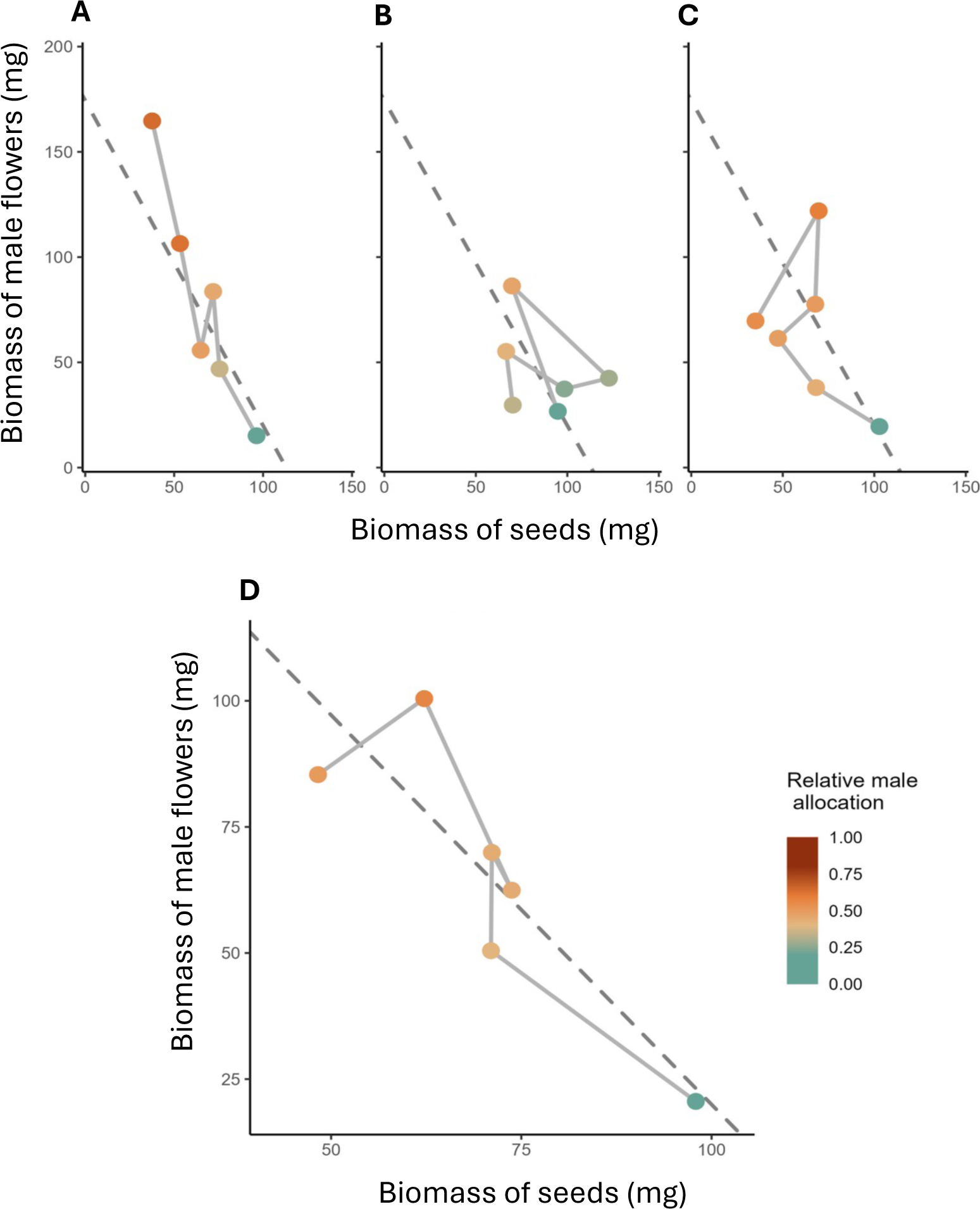
Temporal dynamics of the trade-off in sex allocation between the female and male function from experimental all-female populations of *M. annua* over eight generations (labelled generation 1, 3, 5, 6, 7 and 8), showing average data from each of the three replicate X-only populations (A-C) and average values for seed and male flower per generation for all replicate X-only populations pooled (D). The dotted line shows the result of the standardised major axis regression (intercept = 174, 95%CI [157; 191], slope = -1.54, 95%CI [-1.72; -1.39], R2 = 0.05). The colour of the points reflects the relative male allocation average, calculated as male flower biomass divided by the sum of the biomass of male flowers and seeds together.

**Table 1.** Results of the zero-inflated Gamma generalised linear model, where the response variable is male flower biomass, and the fixed factors are seed mass, plant height, plant slenderness and generations. The total number of individuals retained for this analysis is N=318, with an average of 18 individuals per generation and per population (min = 7, max = 40). The random factors (population, block and phenotyping date) explain 15% of the variance, in addition to the 13% explained by the fixed factors, leading to conditional R^2^=0.28. The calculation of generalized variance-inflation factors (GVIF; Fox and Monette, 1992) shows little multicolinearity between the fixed factors.

|  | GVIF | Coefficient | Standard Error | Confidence interval |
| --- | --- | --- | --- | --- |
| **Intercept** |  | -3.13 | 0.21 | [-3.54, -2.72] |
| **Generations** | 1.08 | 0.08 | 0.04 | [ 0.01, 0.15] |
| **Vegetative traits** |  |  |  |  |
| Height | 1.06 | -0.05 | 0.05 | [-0.16, 0.05] |
| Slenderness | 1.07 | -0.14 | 0.05 | [-0.23, -0.05] |
| **Reproductive traits** |  |  |  |  |
| Seed mass | 1.10 | -0.20 | 0.05 | [-0.30, -0.10] |

### QTL analysis for variation in sex allocation

Allocation to male and female reproductive effort in the *F*_2_ offspring differed markedly among the four crosses (Figure S1). Nevertheless, QTL analyses identified loci associated with variation in sex expression in all four crosses, with the number and genomic location of QTL and the proportion of variance explained differing among the crosses (Figure 3, Table S4). Furthermore, we identified several regions showing strong segregation distortion in crosses 1 (Figure S6), 3 (Figure S8) and 4 (Figure S9). We interpreted these as results of recessive deleterious variants expressed in selfed *F*_2_ crosses and we did not consider QTL overlapping these regions for further interpretation of the results.

**Figure 3.**
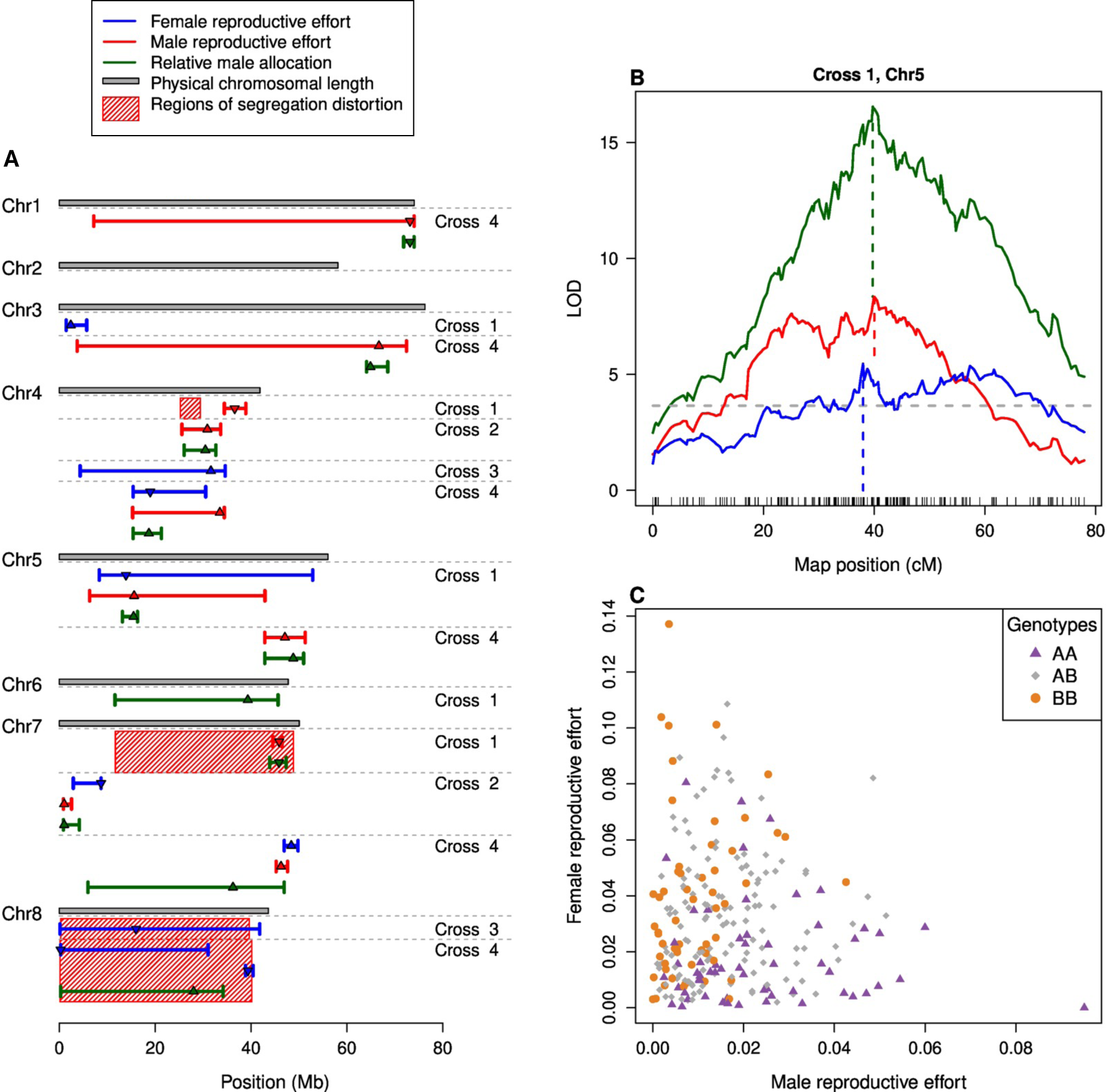
(A) Results of the QTL analyses for from four crosses and their genomic locations three different traits: male reproductive effort (MRE, red bars), female reproducdtive effort (FRE, blue bars), and relative male allocation (rMA green bars). Triangles indicate QTL locations: those pointing up indicate an increase of the trait for alleles inherited from the selection line, while those pointing down indicate a decrease. The length of the bars represents the size of Bayesian credible intervals. Shaded red areas indicate regions of strong segregation distortion on chromosomes in which QTL were located in the same cross. (B) Example from single-QTL model analyses for QTL representing a trade-off locus on chromosome 5 in Cross 1. Dashed vertical lines indicate the location of QTL for MRE (red), FRE (blue), and rMA (green). Vertical black bars along the x axis indicate the location of RAD loci used to build the linkage map. The dashed gray horizontal line designates the significance threshold, inferred using 1000 permutations. (C) Distribution of traits and associated genotypes at the QTL for rMA on chromosome 5 in Cross 1. Purple triangles indicate homozygotes with two alleles inherited from the X-only population (AA), smaller grey diamonds indicate heterozygotes (AB) and orange circles indicate homozygotes with two alleles inherited from the Control population (BB).

Our analysis identified two types loci: isolated small-effect loci, with limited effects on male and female RE in either direction; and loci with a large effect on male sex allocation. For the second type, two loci with antagonistic effects on male and female reproductive effort were identified on the same linkage group. In all cases, for one of these loci, alleles inherited from an individual sampled from a X-only population had a positive effect on male reproductive effort, while, at the other locus, the allele inherited from the X-only population had a negative effect on female reproductive effort. The confidence intervals of both loci overlapped with each other as well as with the inferred confidence intervals of large effect loci for male sex allocation (though the overlap with FRE on chromosome 7 for Cross 2 was with rRA rather than MRE). We thus treat such pairs of ‘loci’ as single trade-off QTL. These trade-off QTL were found on chromosome 5 for Cross 1, on chromosome 7 for Cross 2 and on chromosome 4 for Cross 4 (explaining 24.1%, 32.0% and 8.03% of variance in male sex allocation, respectively). In addition, we identified a second large-effect QTL on chromosome 5 for Cross 4 (explaining 18.6 % of variance in male sex allocation), with a similar pattern of increased male allocation and decreased female allocation, though the QTL for female reproductive effort was just below the *p* < 0.05 threshold of statistical significance in the single-QTL model (Fig S5).

## Discussion

### Implications of phenotypic evidence for a sex-allocation trade-off

Our study has revealed clear evidence for a sex-allocation trade-off in the populations of our selection experiment. A strong phenotypic trade-off between male and female allocations is evident within populations, such that individuals allocating more to their male function (the production of male flowers) allocate correspondingly less to their female function (the production of female flowers and fruits). This inverse linear relationship between male and female allocations is precisely that assumed in theoretical models of sex allocation.

Despite the fundamental nature of the trade-off assumption for sex-allocation theory, it has been difficult to demonstrate the trade-off empirically. As reviewed in the Introduction, many studies have found no relationship between allocations to male and female functions, and others have even found a positive relationship. Those that have been able to demonstrate a trade-off have largely been based on comparisons between individuals with qualitatively different phenotypes (e.g., females versus hermaphrodites), or at the within-individual level (e.g., among flowers). In an artificial selection experiment, Mazer et al. (2007) found quantitative evidence for a trade-off among individuals for a subset of their experimental populations, which evolved lower ovule production when selected for increased pollen production, and vice versa – but other populations showed no such correlated responses to selection. Our study stands out by showing the shape of the expected trade-off among individuals expressing their phenotypes across a broad range of allocations under a controlled by a semi-natural population. It is clear that, in this population, individuals can only produce more pollen if they produce fewer seeds, as assumed by theory.

The successful detection of a sex-allocation trade-off in our study may in part be attributed to the unusually high variation in sex allocation among individuals in our X-only populations. Such high trait variation is often not present in wild hermaphroditic populations in which stabilizing selection is expected to favour a single, canalized sex-allocation strategy. Previous studies have generated variation in sex allocation among individuals by physically removing sexual organs from a proportion of individuals in a population (e.g., Harris and Pannell 2008) or by manipulating organismal physiology (Devisser et al. 1994). While the variation in our study populations was generated under experimental evolution and does not reflect that seen in the wild, the phenotypes generated are nevertheless the outcome of responses to natural selection and the natural expression of genes that affect all aspects of the species’ biology. Our study is thus likely an improvement on more invasive approaches that may interfere with organismal integrity.

Importantly, our experiment has brought about a transition from a dioecious to monoecy. While monoecy is not found in natural populations of diploid *M. annua*, which is uniformly dioecious, it is common in populations with higher ploidy levels, and it is plausible that the transition to monoecy in our experiment illustrates how the natural transition might have occurred in nature in polyploid lineages. In this sense, our study is directly relevant to sex-allocation theory developed to understand the evolution and maintenance of hermaphroditism versus dioecy in general, including transitions involving the gradual evolution of sex allocation in hermaphrodites that accompany the spread of male-and/or female sterility mutations (Charnov et al. 1976, Charlesworth and Charlesworth 1981).

### A quantitative genetic basis of the trade-off in sex-allocation

Our study revealed a shift in the mean sex allocation in the study populations over time, with individuals evolving on average towards greater male allocation and lower female allocation. This shift points to the existence of strong narrow-sense negative genetic covariance between male and female allocations, such that a response to selection on one will be accompanied by an opposite response in the other. Previous work on the genetic basis of male-female correlations has tended to find evidence for broad-sense covariances, which include dominance and epistasis effects that do not link responses to selection directly, and those narrow-sense covariances measured have tended to be positive rather than negative (reviewed in Ashman 2003). The artificial selection experiment of Mazer et al. (2007) showed that selection on either pollen or ovule production study led to negative co-evolution of allocation to the opposite sex, as seen in our experiment. In contrast to the results of Mazer et al. (2007), however, the shift we observed along the male/female trade-off line was almost certainly a response to natural selection principally on male allocation rather than on female function (Cossard et al. 2021).

### QTL and the genetic architecture of the sex-allocation trade-off

Our QTL analyses of all four crosses identified evidence for the segregation of genetic variants associated with phenotypic variation in sex allocation. A particularly remarkable feature of these results is the identification of several ‘trade-off’ loci, which are genomic regions in which alleles inherited from X-only populations cause increased male allocation and decreased female allocation, and vice versa for alleles inherited from Control populations. There has been little research on the genetic architecture of sex allocation, and those studies that have been published have focused almost exclusively on the genetics of sex-ratio variation in dioecious species (Toro and Charlesworth 1982, Carvalho et al. 1998, Vandeputte et al. 2007, Song and Zhang 2024), with Pannebakker et al. (2011) finding evidence for sex-ratio QTL in the dioecious wasp *Nasonia vitripennis*. Spigler et al. (2011) found QTL association with measures of sexual dimorphism in subdioecious strawberry populations and identified a QTL with opposite effects on components of male and female allocation. To our knowledge, the present study is the first to have identified genomic regions in hermaphrodites within which alternative haplotypes or alleles affect sex allocation in opposites directions, as expected for a genetic allocation trade-off assumed by theory (Charnov et al. 1976, Charnov 1982, Charlesworth 1999).

The QTL peaks corresponding to trade-off loci are broad and likely include a great many genes, potentially influencing sex allocation in different ways. First, the QTL peaks may contain single genes with direct antagonistic effects on male and female sex expression. Such genes might affect higher level genetic pathways, e.g., by influencing whether a particular axillary meristem develops as a male or female inflorescences (Cossard et al. 2021). Previous studies have demonstrated that exogenous application of cytokinins can cause changes in sex expression in male and female plants (Li et al. 2019), suggesting that endogenous plant hormone concentrations may directly mediate sex-allocation trade-offs at the point of meristem development. Second, the broad QTL peaks may contain multiple genes with antagonistic effects on either male of female function. While the QTL identified in our study are located in the same chromosomal regions, the distance between peaks for male versus female reproductive effort can be considerable (Figure 3, Table S4). The genomic locations inferred from QTL mapping can be quite precise, despite the inherent limitations of the approach (Price 2006), but multiple loci with effects on sex expression could be co-located in regions with limited recombination, limiting the precision of our QTL analysis. The genome of *M. annua* contains large pericentromeric regions resulting in reduced recombination across substantial parts of all chromosomes (Figure S6-S9). Notably, for the trade-off locus identified on chromosome 5 in Cross 1, the confidence intervals of QTL for both male and female reproductive effort overlap such a pericentromeric region (Figure S6). Such a scenario, where reduced recombination links variants with opposing effects on sex expression, would conform in important respects to the classical model of the evolution of dioecy from hermaphroditism (Charlesworth and Charlesworth 1978) or to the concentration of the effects of multiple loci into a single effective locus (Lesaffre et al. 2024a).

While our study has identified trade-off loci in *M. annua*, with QTL for male and female function co-located in the same chromosomal region with opposite effects, we also identified loci that were associated with variation in allocation to only one of the two sexes. This finding suggests that either genetic variants at certain loci may affect allocation to male function without a direct trade-off against allocation to female function, or that the magnitude of the effects in opposite directions is asymmetric, so that only effects in one direction reach the threshold of statistical significance in our analysis (as in Cross 4 on Chromosome 5, where QTL effects for male and female reproductive effort were in opposite directions, yet with female reproductive effort not significant; see Figure S9). While sex-allocation theory makes the assumption of a direct male-female trade-off, allocation to one sex function may trade-off with allocation to other potential resource sinks such as vegetative growth (Rameau and Gouyon 1991, Seger and Eckhart 1996) – a feature that likely contributes to obscuring sex-allocation trade-offs. In this light, the clear phenotypic and genetic trade-off in sex allocation revealed by our study is all the more striking and points to the overwhelming importance of genomic regions associated with allocation in opposite directions to the two sexual functions.

It is striking that our experiment has seen such strong divergence in sex allocation in response to selection on standing genetic variation. *M. annua* is a colonising species of disturbed habitats, with a metapopulation structure and dynamic (Obbard et al. 2006, Eppley and Pannell 2007, Dorken et al. 2017). In such metapopulations, selection is expected to favour genotypes with a capacity for uniparental reproduction, e.g., hermaphrodites able to produce seed via self-fertilization (Pannell 1997, Pannell and Barrett 1998, Pannell 2015). It is likely that leakiness in sex expression in dioecious *M. annua* is maintained by selection for reproductive assurance during repeated bouts of colonisation of new habitat patches or at low density (Cossard and Pannell 2021), just as monoecy in polyploid populations of the species complex is thought to be maintained by such processes (Obbard et al. 2006, Eppley and Pannell 2007, Dorken et al. 2017). Such dynamics could favour the maintenance of standing variation at a large number of loci with effects on sex expression, of which some will have been sampled in the establishment of our experiment. The strong selection favouring increased male allocation in our X-only populations will then presumably have resulted in an increase in frequency of these variants, a build-up of linkage disequilibrium across loci, and more strongly additive effects of variants that are commonly found as heterozygotes at low frequencies. By demonstrating the simultaneous reduction in female allocation and its genetic architecture, our study provides compelling evidence for the existence of sex-allocation trade-off loci and thus for the fundamental assumption of sex-allocation theory.

## Acknowledgements

We thank T. Kawecki, K. Chen and T. Lesaffre for insightful discussion and comments on the manuscript, A. Revel for assistance in growing plants, numerous undergraduate student assistants for help with phenotyping plants, G. Cossard for managing the first generations of the selection experiment, and the Swiss National Science Foundation (grant 310030_185196) and the University of Lausanne for funding.

## Supplementary Materials

### Supplementary methods

#### Crosses

For a QTL mapping approach, we generated *F*_2_ crosses between plants from the X-only and Control populations. We grew our parental plants from seeds collected from plants from the seventh generation of the experiment. Besides regular selection populations, where the next generation grown from pooled seeds from all individuals in the population, additional artificial selection populations have been established in the previous generation. Plants from the artificial selection populations were grown from seeds, which were among the top 20% of the pollen producers. The plants in this treatment showed an even stronger increase in pollen production than the regular selection populations (Xinji Li, unpublished results).

As pollen donors, we chose to use plants with high pollen production from either an artificial selection population (Cross 2) or from a regular selection population (Crosses 1, 3 and 4). As seed donors we chose female plants from one of the Control populations, which did not show any leakiness. Seeds were planted in horticultural soil and plants were grown in greenhouses until they started producing inflorescences. A single selected female as pollen donor and multiple females as ovule producers from the Control populations were transferred to closed crossing boxes. In these boxes, plants were left to openly pollinate for several weeks after which seeds and tissue samples for DNA extraction were collected.

We grew *F*_1_ plants from seeds and made crosses the same way as we did with their parents with the difference that only single individuals were transferred to crossing boxes, so that all seeds were the result of selfing. We grew *F*_2_ plants from the resulting seeds in the greenhouse until they reached maturity. We collected, dried and weighed all male (*W_m_*) and female flowers including seeds (*W_f_*) and also collected, dried and weighed the remaining plant material (*W_p_*). However, after phenotyping the first plants from Cross 1 we realized that this phenotyping protocol would be too time intense and we changed the protocol and collected male and female flowers on every second branch. For these samples we multiplied *W_m_* and *W_f_* times two in the denominator of the calculations of MRE and FRE. In addition, we realized that *F*_2_ plants in Crosses 3 and 4 were pollen limited, so that not all female flowers were fertilized and the dry weight of female flowers would underestimate female sex allocation. To overcome this issue, we counted the number of female flowers (fertilized or unfertilized) on every second branch in these two crosses. We then estimated the dry weight of flowers based on 30 individuals for which we both counted and weighed dried female flowers. In addition we collected tissue samples, extracted DNA using a TECAN extraction robot and quantified it using a Cybr Green protocol on a TECAN Plate reader.

For each plant we computed male and female allocations in terms of male (MRE) and female reproductive effort (FRE), respectively, defined as the proportion of reproductive biomass (male flowers for MRE and female flowers, fruits and seeds for FRE) relative to total plant biomass. Thus, MRE = *W*_m_/(*W*_m_ + *W*_f_ + *W*_p_) and FRE = *W*_f_/(*W*_m_ + *W*_f_ + *W*_p_). We also computed an index for relative male sex allocation, rMA = MRE/(MRE + FRE).

#### Genomic analysis

We generated genotype data using a modified ddRad protocol (Peterson et al. 2012). For each sample we first digested about 100 ng DNA diluted to 22 μl by adding 2.5 μl Smartcut buffer (NEB), 0.4 μl EcoRI-HF (NEB) and 0.4 μl Taq1 (NEB) restriction enzymes and incubating at 37 °C for 30 min, 65 °C for 30 min and 80 °C for 20 min. We then ligated P1 and P2 adapters described in Peterson et al. (2012), which were diluted to 40 μM in 1x annealing buffer (50mM NaCl, 10 mM). We added 3 μl rATP (10 mM), 2 μl annealed P2 adapter (3 μM), 0.8 μl 10x T4 Ligase buffer (NEB) 1 μl T4 Ligase (400 U/μl, NEB), 22 μl digested DNA and 2 μl annealed P1 adapter (0.3 μM) and incubated for 20 min at 23 °C followed by 10 min at 65 °C. We then size selected 300 μl of pooled samples for each library to 550 bp. For this we first added 0.57x volume of Ampure XP beads incubated for 10 min at room temperature and saved the supernatant, to which we added 0.12x volume of Ampure XP beads (Beckman Coulter), incubated on a magnetic stand, washed the beads with 70% ethanol and eluted DNA in 30 μl water for 2 min.

We selected for biotin-labeled P2 adapters using M-270 Dynabeads (Invitrogen). We washed 15 μl Dynabeads 3 times in 1x bind and wash buffer (5 mM Tris-HCl pH 7.5, 0.5 mM EDTA, 1M NaCl), resuspended them in 30 μl 2x bind and wash buffer, added the size selected DNA and incubated for 15 min at room temperature. We discarded the supernatant, washed the beads three times using 1x bind and wash buffer and resuspended the beads in 45 μl water. We then amplified size selected fragments by PCR, using dual indexed primer pairs. We added 45 μl bead suspension, 3 μl forward primer (10 μM), 3 μl reverse primer (10 μM) and 50 μl KAPA HiFi Hotstart Ready Mix (Roche). PCR program was 2 min at 95 °C initial denaturation and 11 cycles at 98 °C for 20 s, 65 °C for 20 s and 72 °C for 30 s.

We cleaned PCR products in 0.7x Ampure XP beads according to manufacturer’s instructions and eluted PCR products in 20 μl water. Cleaned libraries were sequenced using 150 bp paired-end reads on the Illumina Novaseq S6 platform by Novogene UK. We demultiplexed and quality filtered raw Illumina reads using Stacks 2.62 (Catchen et al. 2013). The following part has been integrated in a reproducible Snakemake workflow (Mölder et al. 2021) (https://github.com/jgerchen/qtl_mapping). We aligned reads against the diploid female *M. annua* genome assembly (Christenhusz et al. 2024) using BWA mem 0.7.17-r1188 (Li 2013) and sorted and indexed alignment using samtools 1.15.1 (Danecek et al. 2021). We then used the gstacks module in Stacks for reference-based variant calling and used the population module to generate a VCF file for the cross, excluding sites with more than 50% missing data (option -R 0.5). Based on the results of gstacks, we identified individuals with an average sequencing coverage smaller than 10 and removed them using vcftoools 0.1.16 (Danecek et al. 2011).

We then made a Principal Component analysis based on genotypes using R with the packages Adegenet 2.1.10 (Jombart 2008), vcfR 1.15.0 (Knaus and Grünwald 2017) and Pegas 1.3 (Paradis 2010). *F*_2_ offspring from a single cross is expected to cluster at the first two principal components with the *F*_1_ parent, which provided all the genetic variation found in the *F*_2_. In contrast, the *F*_0_ grandparents are expected to be more distant, since they only contributed half of their genetic variation each to the *F*_1_. We manually identified and removed *F*_2_ individuals that did not cluster as expected as putative pollen contaminants. We then ran further filtering using vcftools to remove loci, which had a minor allele frequency smaller than 0.2 and did not retain two alleles after filtering.

We used LepMap3 0.5 (Rastas 2017) to build new linkage maps for each cross. First, we estimated sex-averaged (sexAveraged=1) linkage maps for each Chromosome using the OrderMarkers2 module twice, once with fixed markers order based on chromosomal locations (physical maps, improveOrder=0), and once allowing LepMap to infer a new marker order (genetic maps, improveOrder=1). When building new linkage maps, LepMap places problematic loci at the ends of linkage maps. We used this to filter out loci, by comparing their location on physical and genetic maps. We plotted these and manually removed loci where physical and genetic maps strongly disagreed and which were placed at the ends of genetic maps. We then used the OrderMarkers2 module to phase remaining loci based on grandparent genotypes. Since not all loci are informative for grandparent phase, we ran OrderMarkers2 twice again, with phasing based on grandparent genotypes either turned on or off (option grandParentphase 1 or 0 respectively). We then used the scripts phasematch.awk and map2genotypes.awk, which are included with lepMap3, to impute phase for uninformative loci and generate genotypes based on imputed maps and used a custom python script to transform genotypes to rQTL input format. In addition, we used TRASH 1.2 (Wlodzimierz et al. 2023) to identify centromeric repeats on the *M. annua* genome assembly to better understand the genomic context of our results.

#### QTL analysis

We used the R package rQTL 1.10 (Arends et al. 2010) to do the QTL analysis for each cross. First, we inferred genotype probabilities using the calc.genoprob function with a stepsize of one. We then inferred single QTL models for all three traits (MRE, FRE and rMA) using Haley-Knott regression (Haley and Knott 1992) with the scanone function of rQTL, with 1000 permutations to infer LOD cutoff values. We then fitted two-QTL models using the scantwo function with 10000 permutations to infer LOD cutoff values and tried to identify interactions between loci. Finally, based on the results of the single- and two-QTL models we fitted multi-QTL models. We used the makeqtl function to generate a QTL object, based on the results from the single-QTL analysis. We then used the fitqtl function using to fit the multi-QTL model using Haley-Knott regression. We then used the refineqtl function to refine QTL locations based on the multi-QTL model and rerun the fitqtl function again to get the final fit of the model. Finally, we reran the fitqtl function with the options dropone set to FALSE and get.ests set to TRUE to estimate effect sizes of QTL from the multi-QTL model.

### Supplementary Results

#### Sex allocation in the parental generation for QTL analysis

None of the plants from the Control populations used as maternal grandparents for the crosses produced any male flowers. In contrast, MRE ranged from 0.001 to 0.024 and rMA from 0.007 to 0.96 in the plants used as maternal grandparents from the X-only populations. Resulting *F*_1_ plants produced both male and female inflorescences (Table S1). Note that only *F*_2_ offspring from the same cross were grown in a common garden, so that phenotypes between crosses or between generations within crosses are not directly comparable. No phenotypic information is available for the *F*_0_ from the Control population and the *F*_1_ of Cross 4, because the plants died before they could be phenotyped. However, both plants produced sufficient male flowers to sire either *F*_1_ offspring via outcrossing or *F*_2_ offspring via selfing respectively.

#### Offspring phenotypes for the QTL analysis

We phenotyped and genotyped a total of 257, 376, 374 and 332 *F*_2_ offspring for crosses 1,2,3 and 4 respectively. We removed 7, 2, 21 and 6 Individuals, which had low sequencing coverage and 3, 6, 2 and 4 Individuals, which we identified as pollen contaminant based on the results of PCA analyses on genotypes for Crosses 1, 2, 3 and 4 respectively. In addition we removed two individuals from Cross 3, which were phenotypic outliers for MRE and one individual from Cross 4, which was a phenotypic outlier for FRE (Figure S1).

#### Linkage maps

Final linkage maps had a total length of 605.68 cM, 616.27 cM, 583.6 cM and 649.25 cM and included 2519, 1972, 2573 and 2835 RAD-loci for Crosses 1, 2, 3 and 4 respectively. We found several peaks of strong segregation distortion in Crosses 1, 3 and 4, which were likely the result of inbreeding depression due to selfing of *F*_1_ individuals (Figures S6, S8 and S9).

#### Single-QTL model

Our single-QTL analyses found evidence for QTL in all four crosses. For Cross 1, evidence for QTL for all three traits was found on Lg5. In addition, further QTL for MRE were found on Lg4 and Lg7 as well as additional QTL for MRE on Lg3 and for rMA on Lg6 (Figure S2). For Cross 2, QTL for all three traits were found on Lg7. In addition, QTL for MRE and rMA were found on Lg4 (Figure S3). For Cross 3 QTL were found for MRE on Lg4 and Lg8 (Figure S4). For Cross 4, three QTL were found for FRE on Lg4, Lg7 and Lg8, for MRE five QTL were found on Lg1, Lg3, Lg4, Lg5 and Lg7 and for rMA six QTL were found on Lg1, Lg3, Lg4, Lg5, Lg7 and Lg8 (Figure S5).

#### Two-QTL model

The two-QTL analysis found only two cases in which there was evidence for either additional QTL that were not found using the single-QTL model. In Cross 1, the two-QTL model identified an additional QTL for rMA on Lg7. In Cross 4 it identified a second QTL for MRE on lg8 (Table S2). However, in both cases these additional QTL are also co-localized with strong signatures of segregation distortion (Figures S6 and S9). Since we didn’t consider these QTL trustworthy, we ran the follow-up multi-QTL model twice, once including the additional QTL and other QTL located in areas of strong segregation distortion and once without them.

#### Multi-QTL model

The final results from the multi-QTL models showed that the amount of variation explained by the QTL varied between 5.23% for FRE for Cross 3 and 44.46% for rMA for Cross 4 (47.45% when including QTL located in regions with strong segregation distortion, Table S3). Overall, all individual QTL inferred by the single- and two-QTL model had highly significant statistical support in the multi-QTL model and the proportion of variance explained by individual QTL ranged from 3.19% to 31.99% (Table S3). The distribution of QTL varied strongly between crosses. While the greatest part of the variation was explained by QTL with antagonistic effects on MRE and FRE on the same chromosome (Figures S6 and S7), in Cross 3 only limited variation in FRE was explained by two QTL, one of which was located in an area with strong segregation distortion (Figure S8). In contrast, in Cross 4, a large proportion of the trait variation is explained by multiple QTL of small to medium effect size (two for FRE and five for MRE excluding QTL in areas of strong segregation distortion, Figure S9).

**Table S1.**
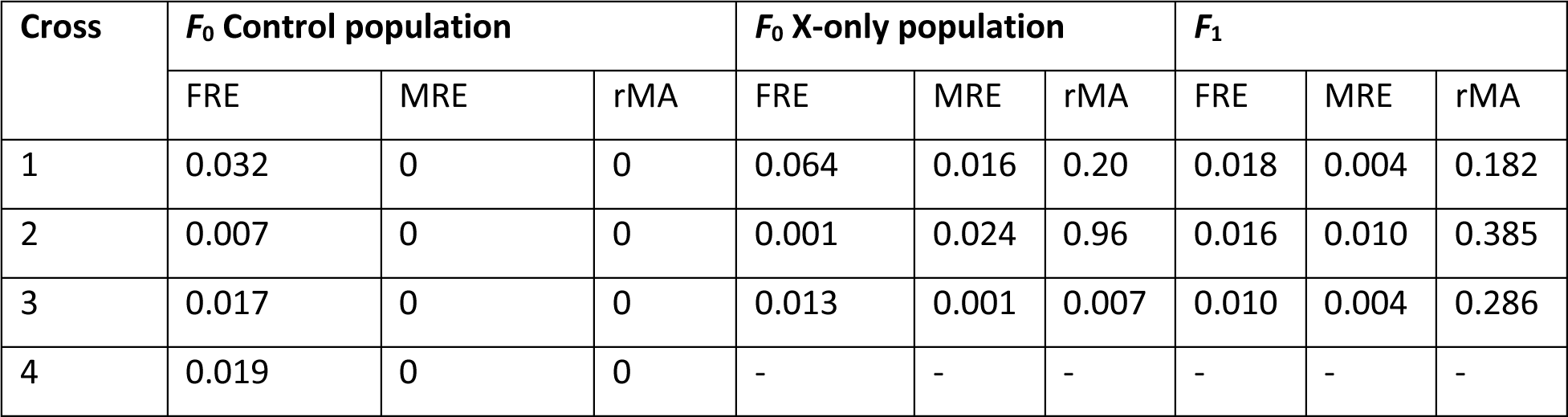
Sexual phenotypes of *F*_0_ grandparents from Control (female parent) and X-only populations (male parent) and selfed *F*_1_ parents of crosses.

**Table S2.** Results from the QTL analysis, which are either indicative of additional QTL not found in the single-QTL analysis or for interactions between QTL. Lg1 and Lg2 are linkage groups on which the two QTL are found, Pos1 f, Pos 2 f and Pos 1 a and Pos 2 a are the positions of QTL based on full and additive models. LOD fv1, LOD int and LOD av1 are LOD scores indicating whether either the full model, an interaction or an additive model are superior to simpler one-QTL models and *p* fv1, *p* int and *p* av1 are the resulting *p*-values. Additional QTL and statistically significant *p*-values are highlighted in bold. Complete results can be found in supplementary file 1.

| Cross | Lg1 | Lg2 | Pos1 f<br>(cM) | Pos2<br>f<br>(cM) | LOD<br>fv1 | $p$ fv1 | LOD<br>int | $p$ int | Pos1a<br>(cM) | Pos2a<br>(cM) | LOD<br>av1 | $p$ av1 |
| --- | --- | --- | --- | --- | --- | --- | --- | --- | --- | --- | --- | --- |
| 1 | 5 | **7** | 40 | 62 | 5.194 | 0.724 | 0.431 | 1 | 40 | 59 | 21.21 | **0.003** |
| 4 | 8 | **8** | 1 | 30 | 8.53 | 0.283 | 1.088 | 1 | 1 | 32 | 7.451 | **0.016** |

**Table S3.** Results of the multi-QTL analyses for the full model. Traits that end with ‘sd’ include also QTL that were found in regions with strong evidence of segregation distortion. % var refers to the proportion of variance explained by the full model.

| Cross | Trait | % var | *p* (F) |
| --- | --- | --- | --- |
| 1 | FRE | 17.56 | 1.58e-09 |
| 1 | MRE | 24.92 | 2.69e-14 |
| 1 | MREsd | 35.32 | 0 |
| 1 | rMA<sub>sd</sub> | 35.75 | 0 |
| 2 | FRE | 20.78 | 0 |
| 2 | MRE | 35.93 | 0 |
| 2 | rMA | 37.03 | 0 |
| 3 | FRE | 5.23 | 9.27e-05 |
| 3 | FREsd | 9.7 | 4.25e-07 |
| 4 | FRE | 13.92 | 1.19e-09 |
| 4 | FREsd | 31.93 | 0 |
| 4 | MRE | 38.09 | 0 |
| 4 | rMA | 44.46 | 0 |
| 4 | rMA<sub>sd</sub> | 47.45 | 0 |

**Table S4.**
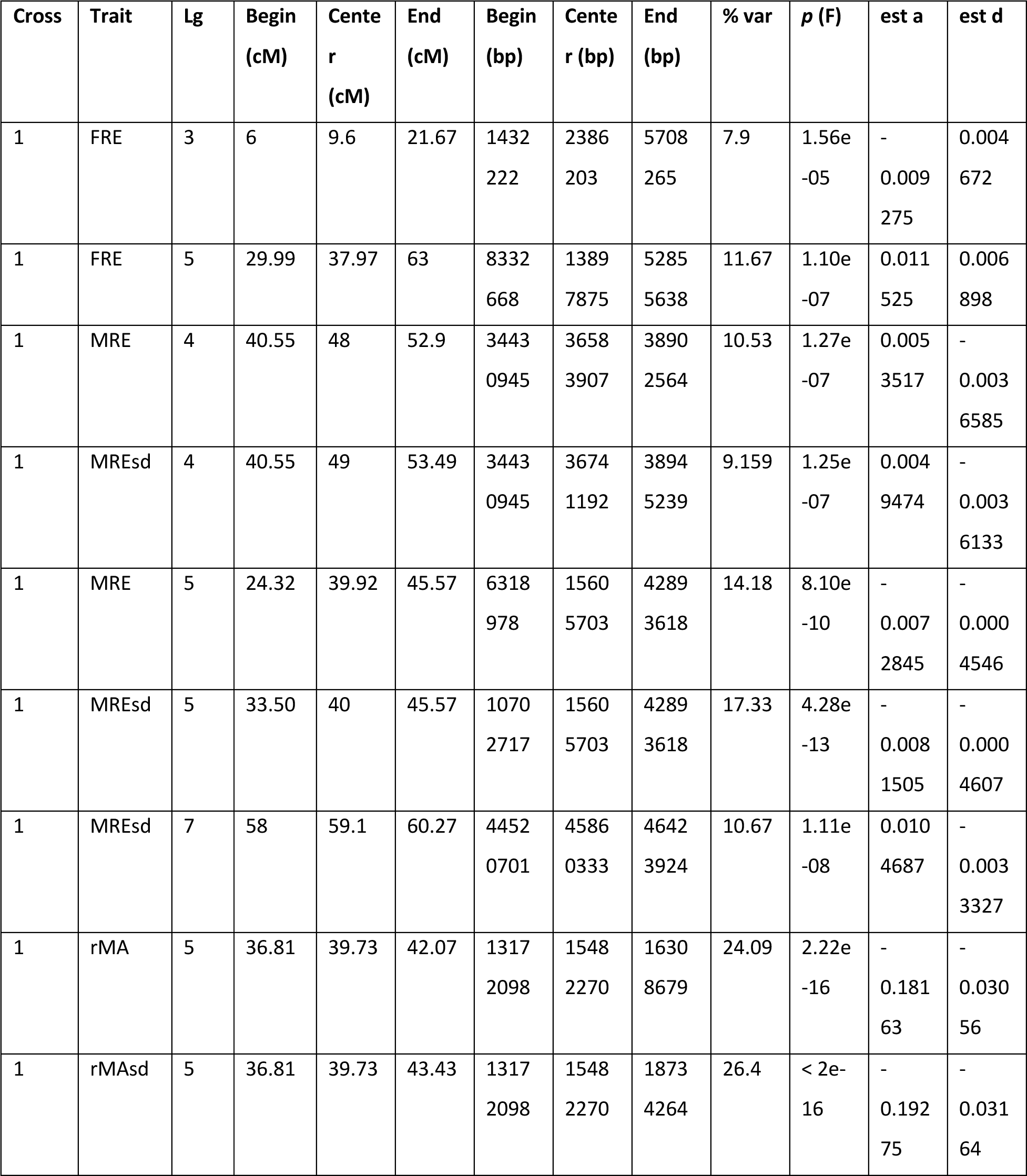
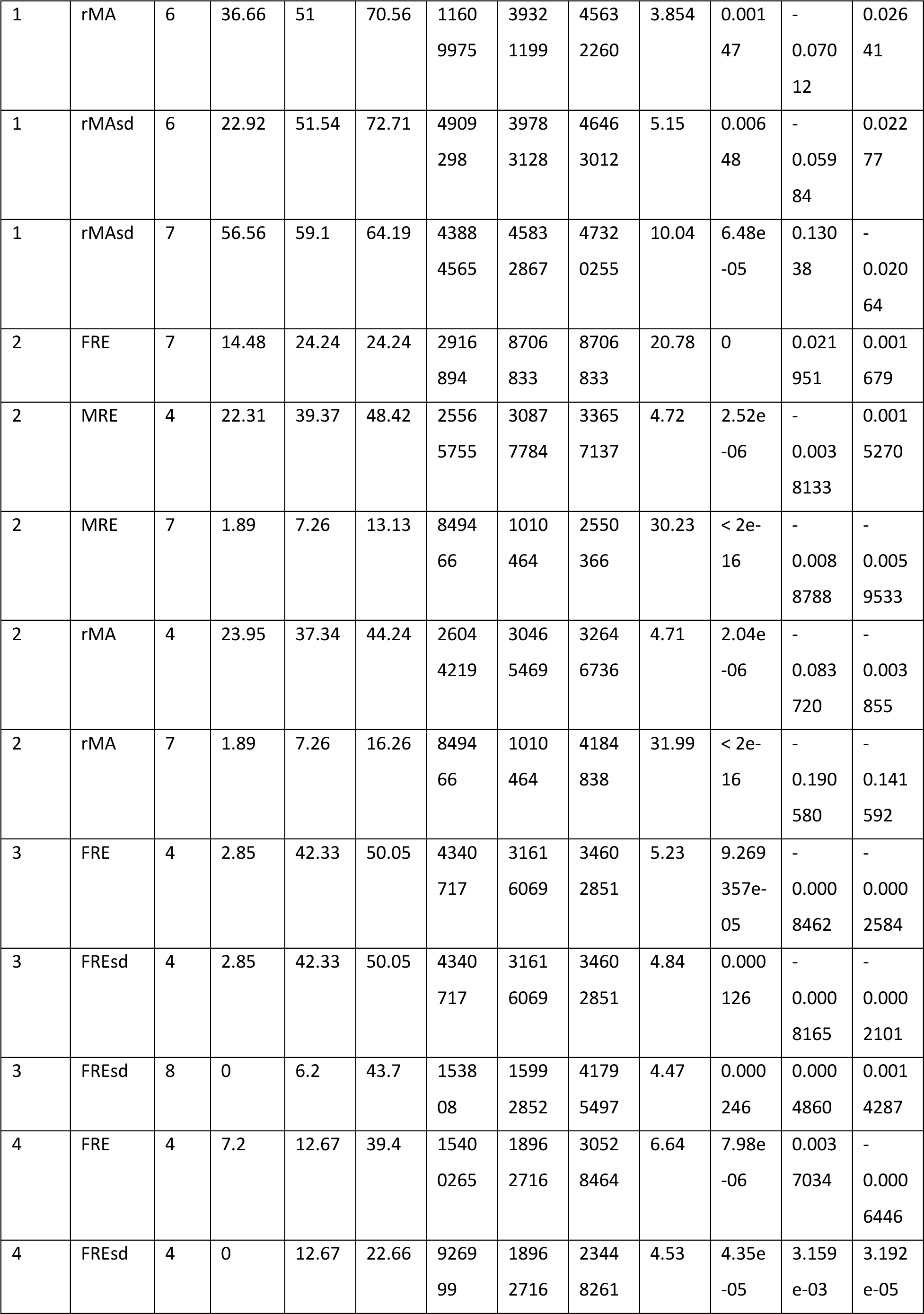
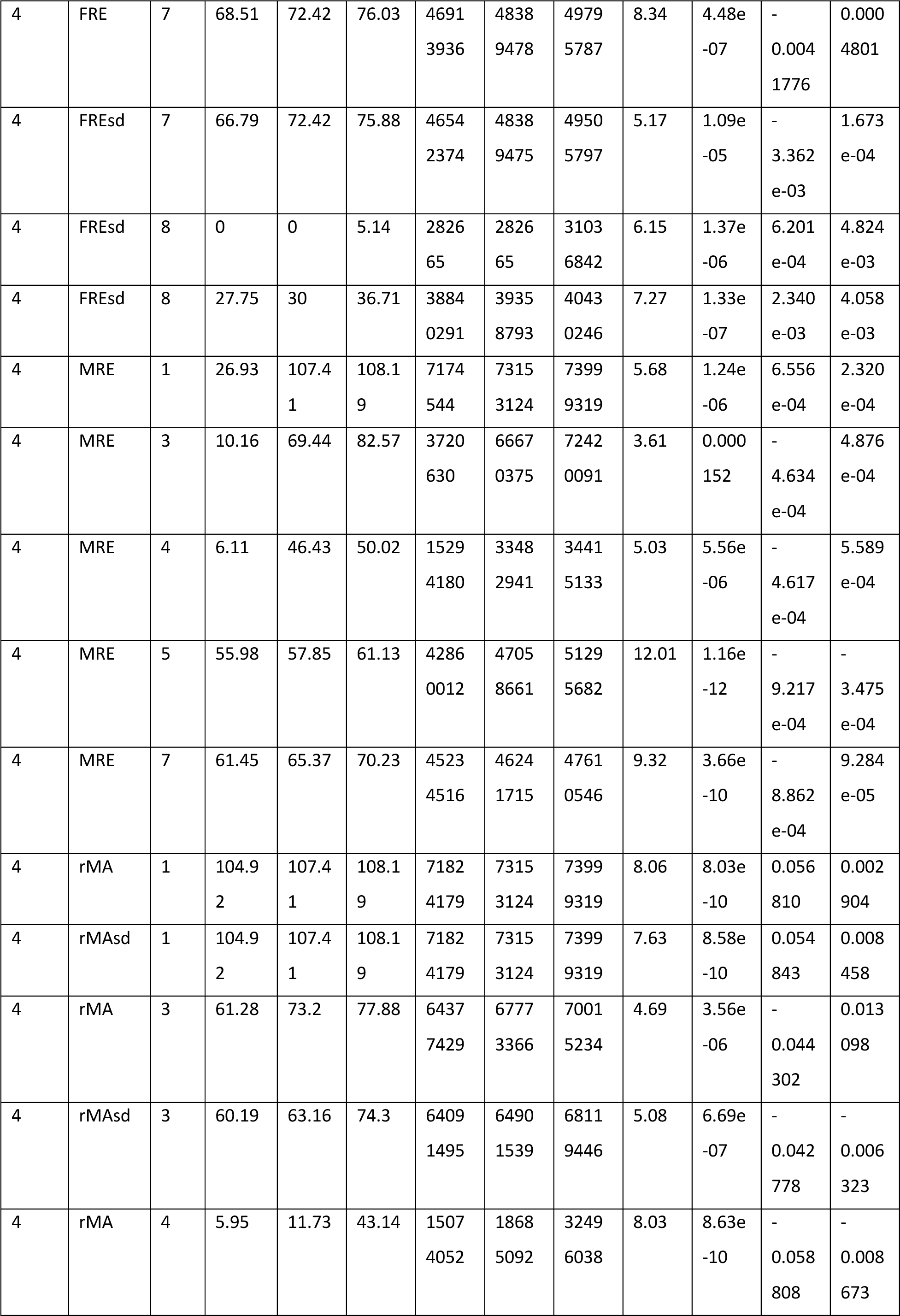
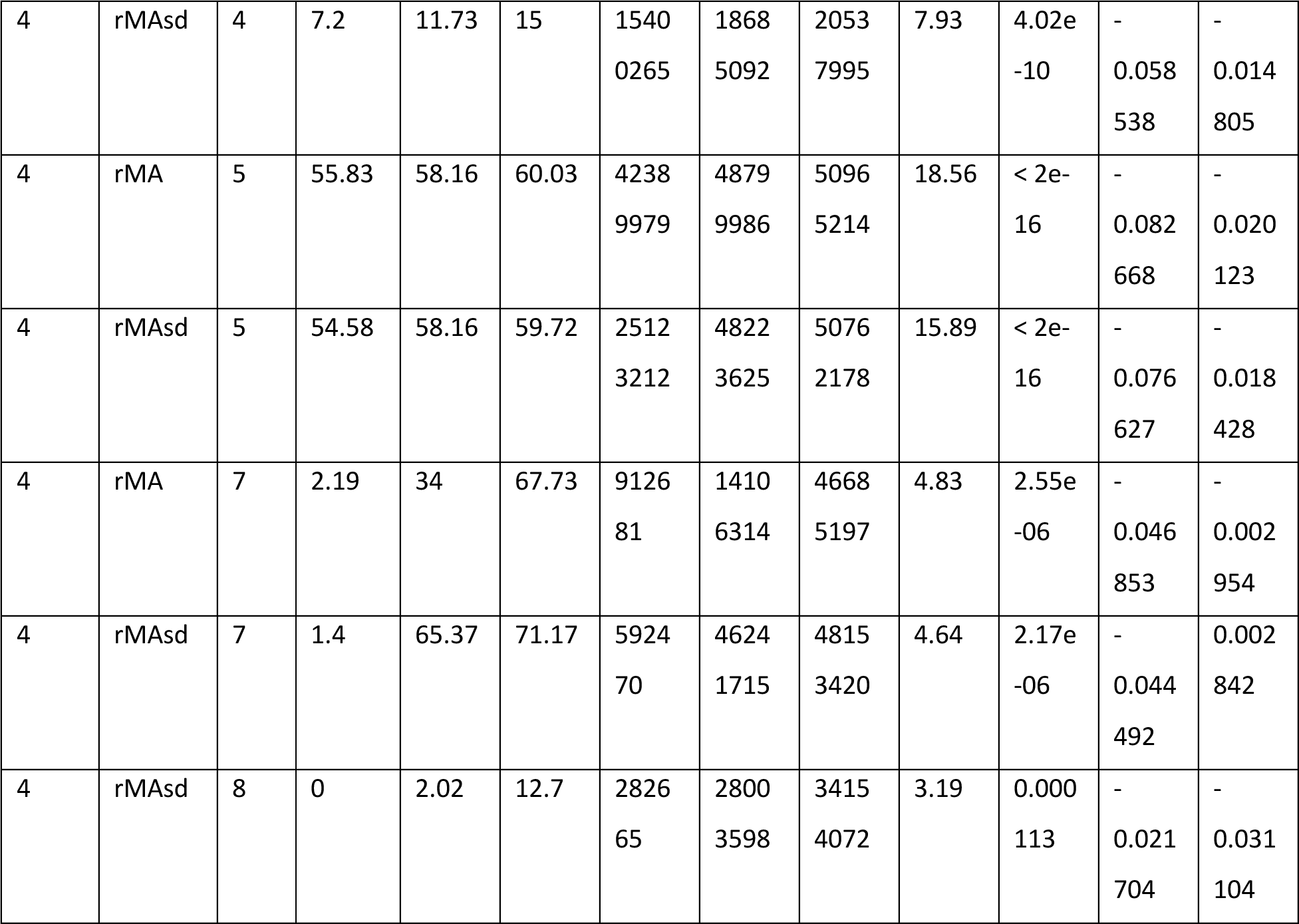
Results of the multi-QTL analyses for individual QTL. Traits that end with ‘sd’ include also QTL that were found in regions with strong evidence of segregation distortion. Begin and End indicate the limits of Bayesian confidence intervals in either cM or bp. % var refers to the proportion of variance explained by the single QTL. *p* (F) is the p-value from an ANOVA when dropping the single QTL. est a and est d are the estimated additive and dominance effects of the QTL.

**Figure S1.**
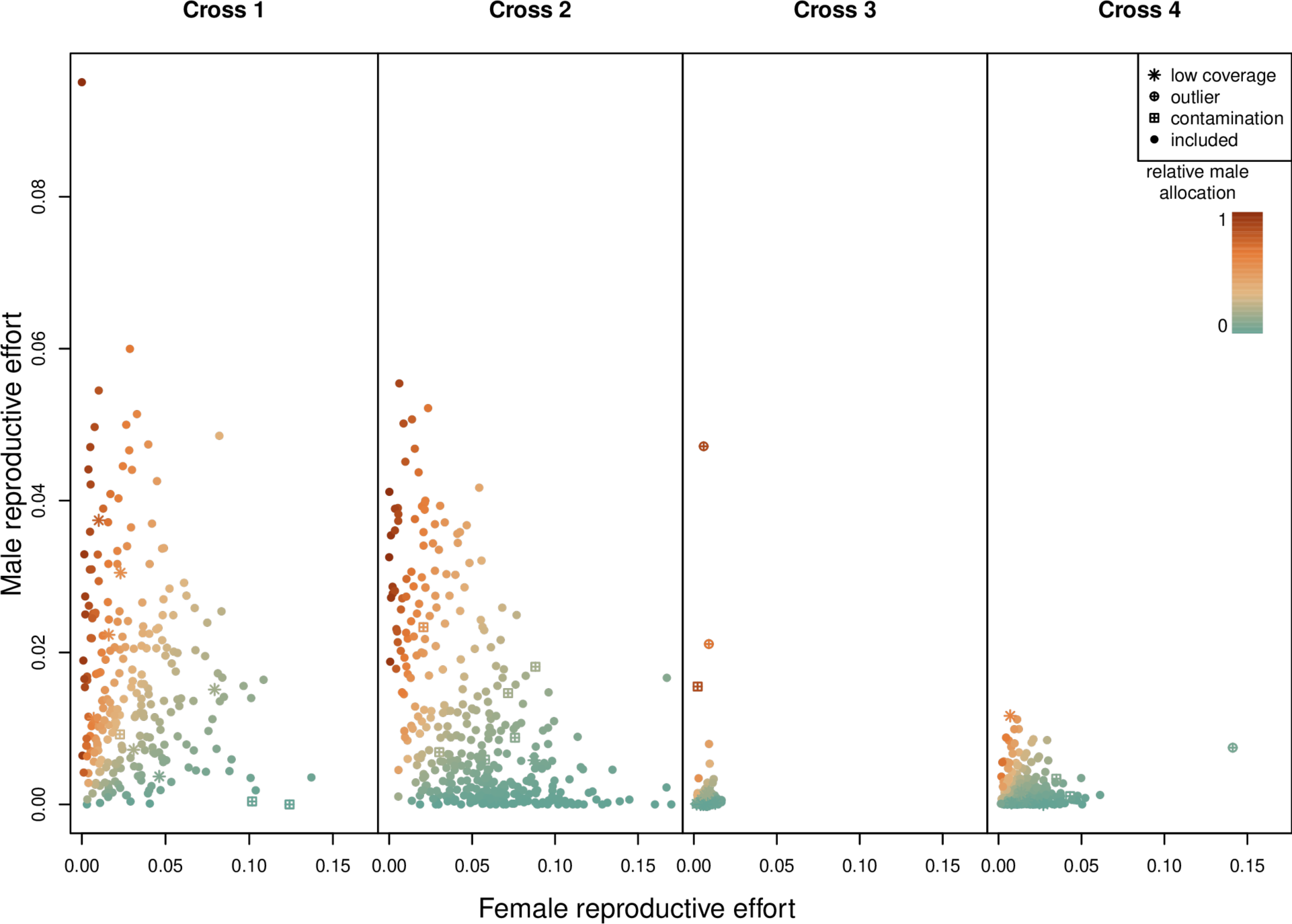
Female reproductive effort (FRE) and male reproductive effort (MRE) for *F*_2_ offspring of all four crosses. Colour coding indicates relative male allocation (rMA). Icons indicate samples that were removed from the final QTL analysis which either had low sequencing coverage, were phenotypic outliers, or were inferred to be the result of pollen contamination based on principle component analysis of genotypes.

**Figure S2.**
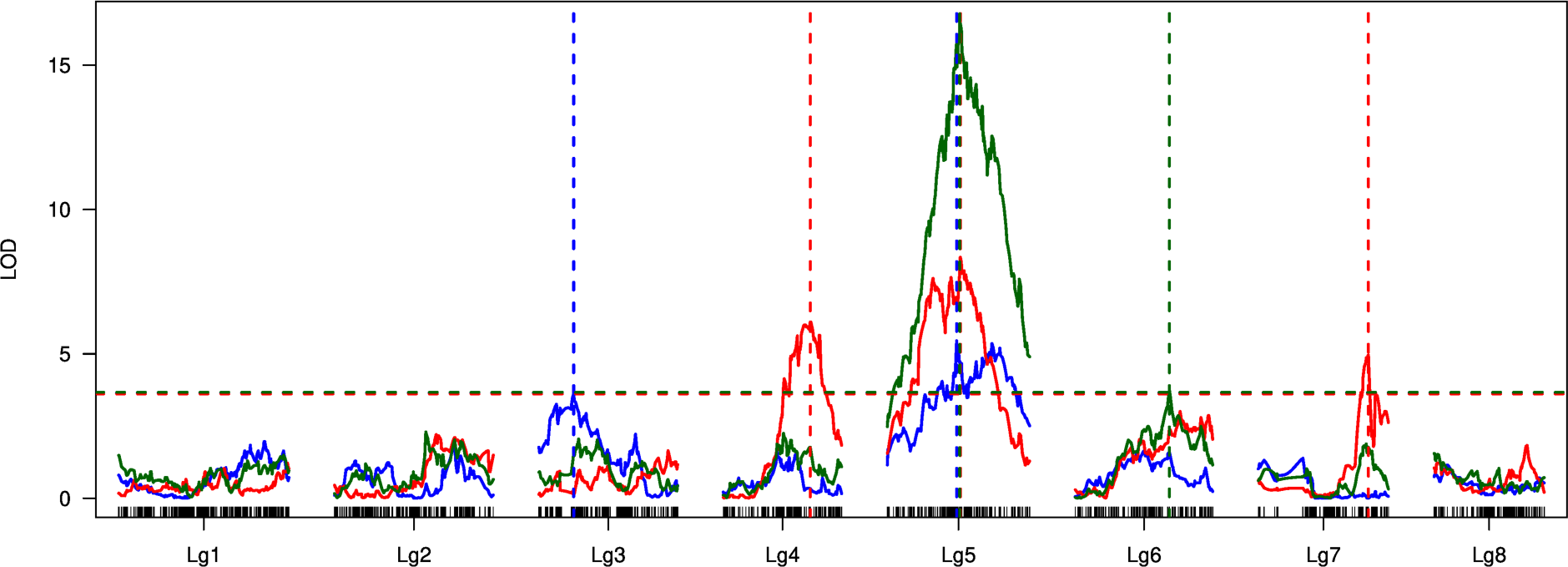
Result from single-QTL analyses for Cross 1. Colours indicate FRE (blue), MRE (red) and rMA (green). Dashed horizontal lines indicate significance thresholds (*p* = 0.05) inferred by 1000 permutations, vertical dashed lines indicated positions of QTL. Black bars at the bottom indicate location of RAD loci on linkage maps.

**Figure S3.**
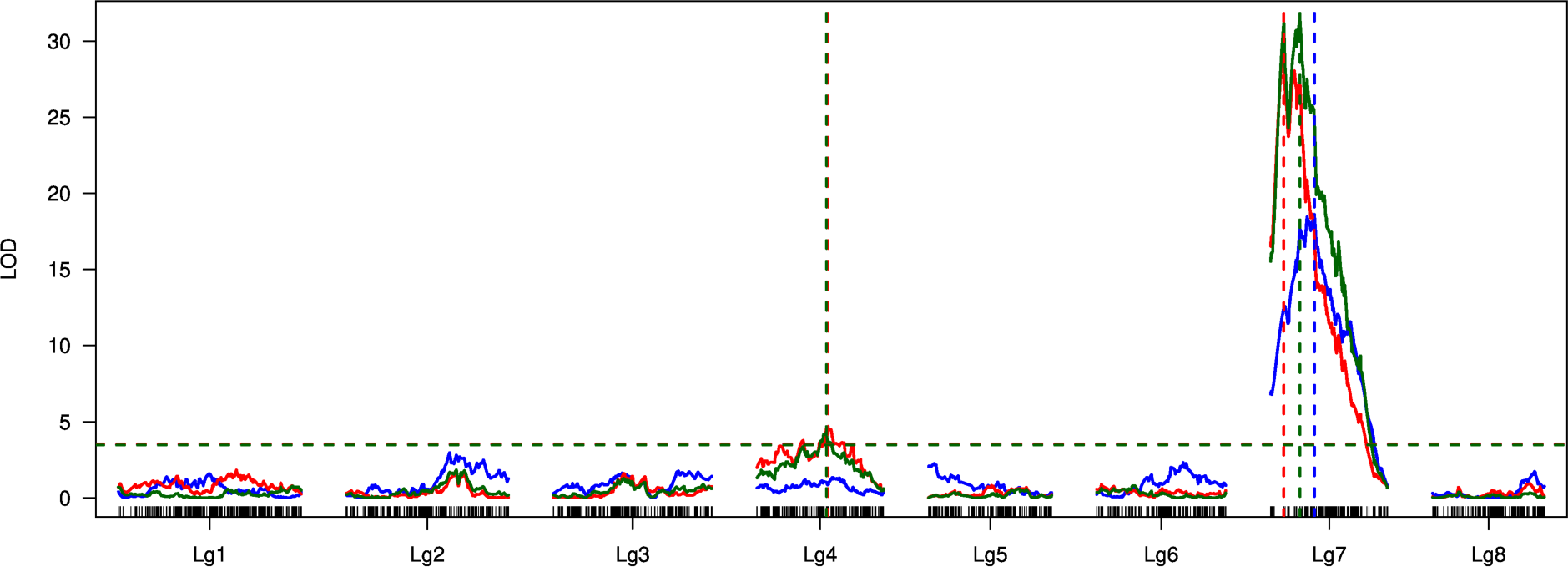
Result from single-QTL analyses for Cross 2. Colours indicate FRE (blue), MRE (red) and rMA (green). Dashed horizontal lines indicate significance thresholds (*p* = 0.05) inferred by 1000 permutations, vertical dashed lines indicated positions of QTL. Black bars at the bottom indicate location of RAD loci on linkage maps.

**Figure S4.**
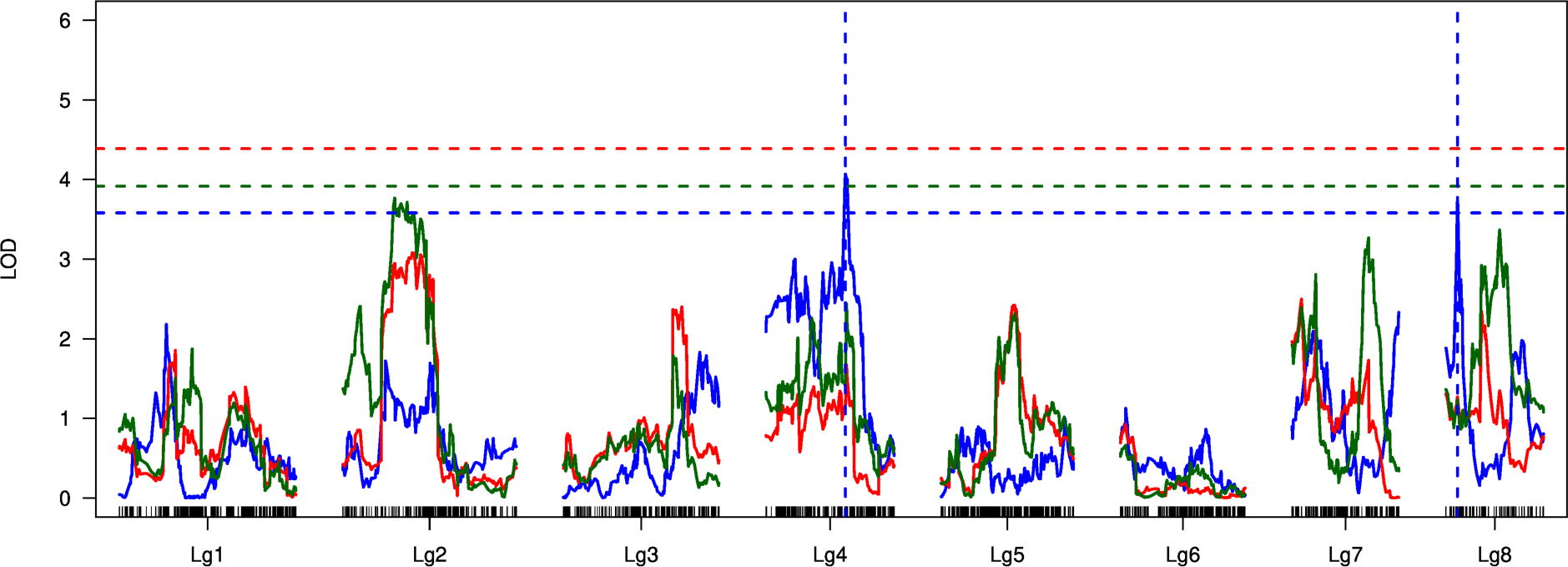
Result from single-QTL analyses for Cross 3. Colours indicate FRE (blue), MRE (red) and rMA (green). Dashed horizontal lines indicate significance thresholds (*p* = 0.05) inferred by 1000 permutations, vertical dashed lines indicated positions of QTL. Black bars at the bottom indicate location of RAD loci on linkage maps.

**Figure S5.**
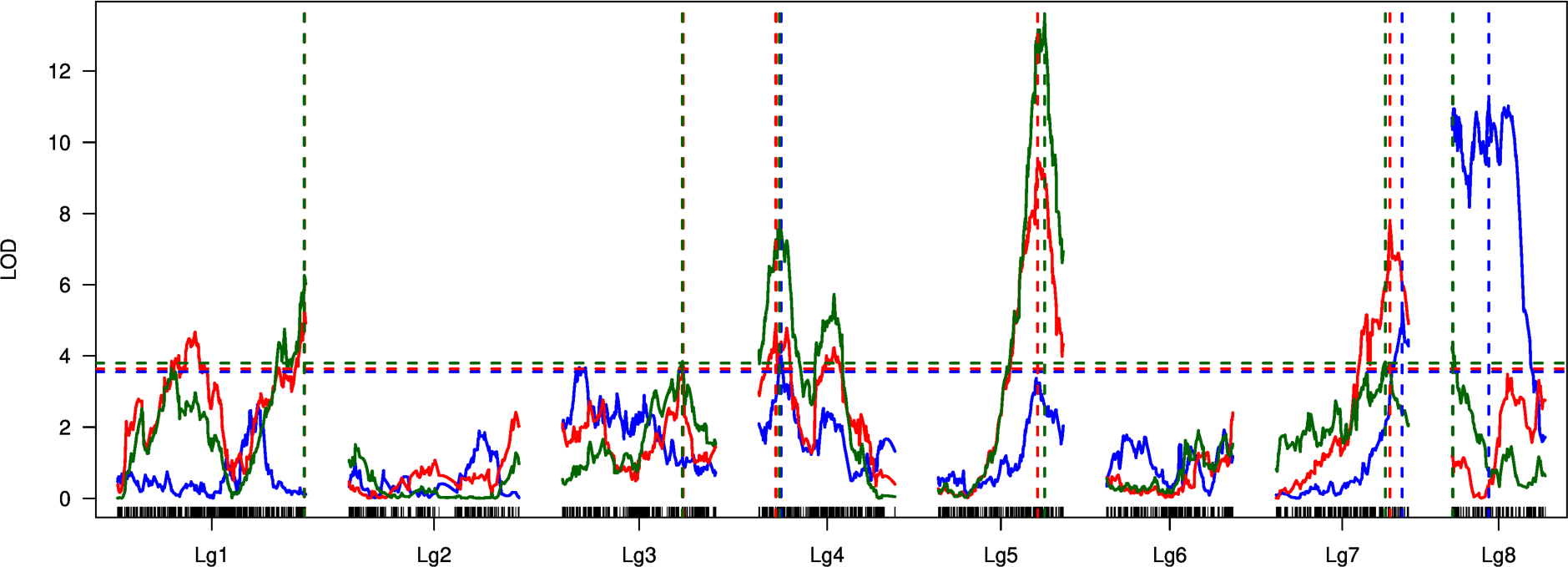
Result from single-QTL analyses for Cross 4. Colours indicate FRE (blue), MRE (red) and rMA (green). Dashed horizontal lines indicate significance thresholds (*p* = 0.05) inferred by permutations, vertical dashed lines indicated positions of QTL. Black bars at the bottom indicate location of RAD loci on linkage maps.

**Figure S6.**
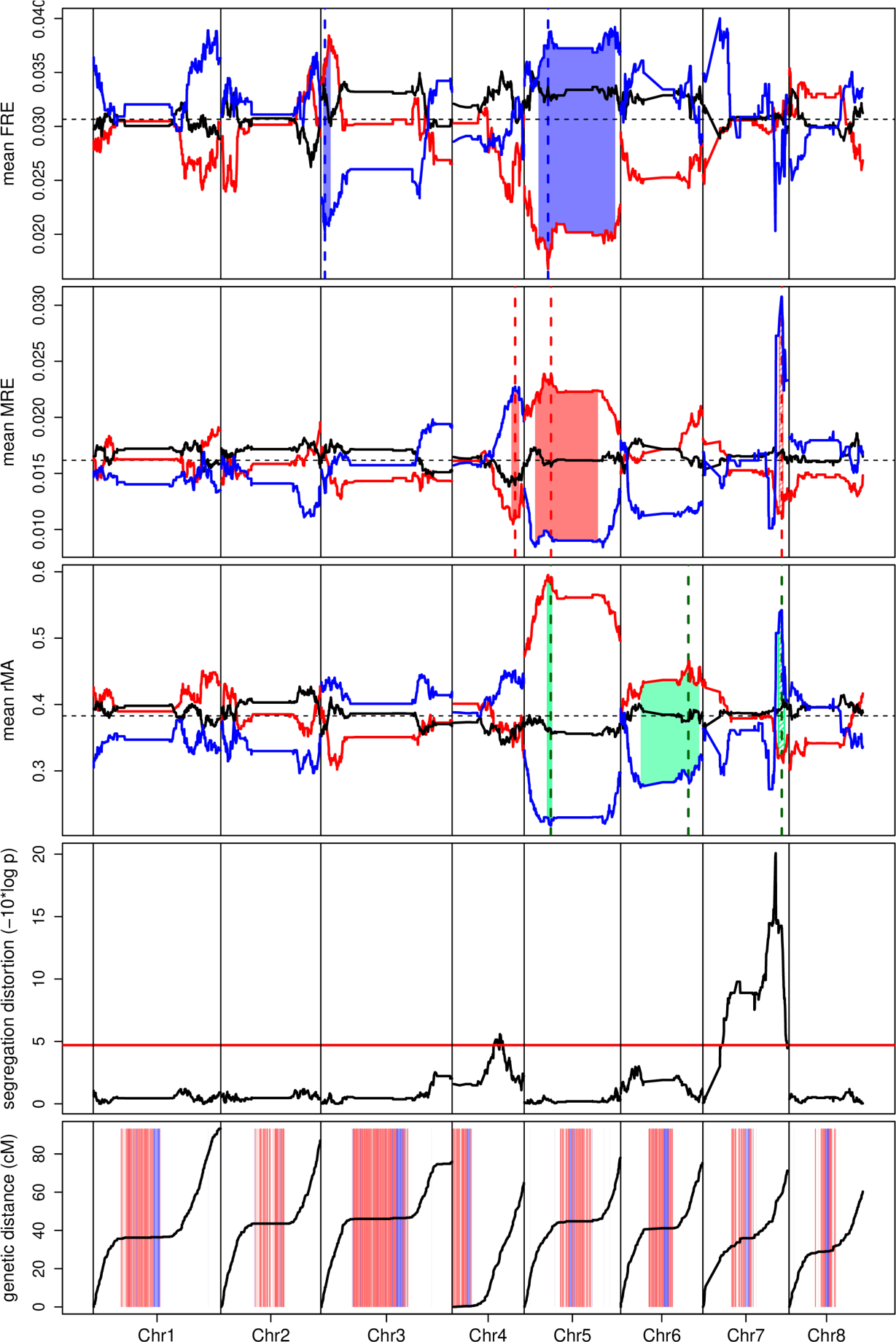
Genomic results for Cross 1. Top three plots: Mean traits (FRE, MRE, rMA) across the genome. Thick red, blue and black lines indicate mean trait values for homozygotes inherited from either Control (blue) or X-only populations (red) or heterozygotes. The horizontal, dashed line indicates mean trait values for all individuals. Vertical dashed lines indicate the location of QTL, shaded areas indicate the Bayesian credible intervals for QTL. Plot 4: *p*-value of χ²-test for segregation distortion. The red line indicates the significance threshold (*p* < 0.05) after Bonferroni correction. Bottom plot: recombination maps inferred from RADseq data. Shaded red and blue bars in the background indicate genomic windows enriched for the two most common centromeric repeats.

**Figure S7.**
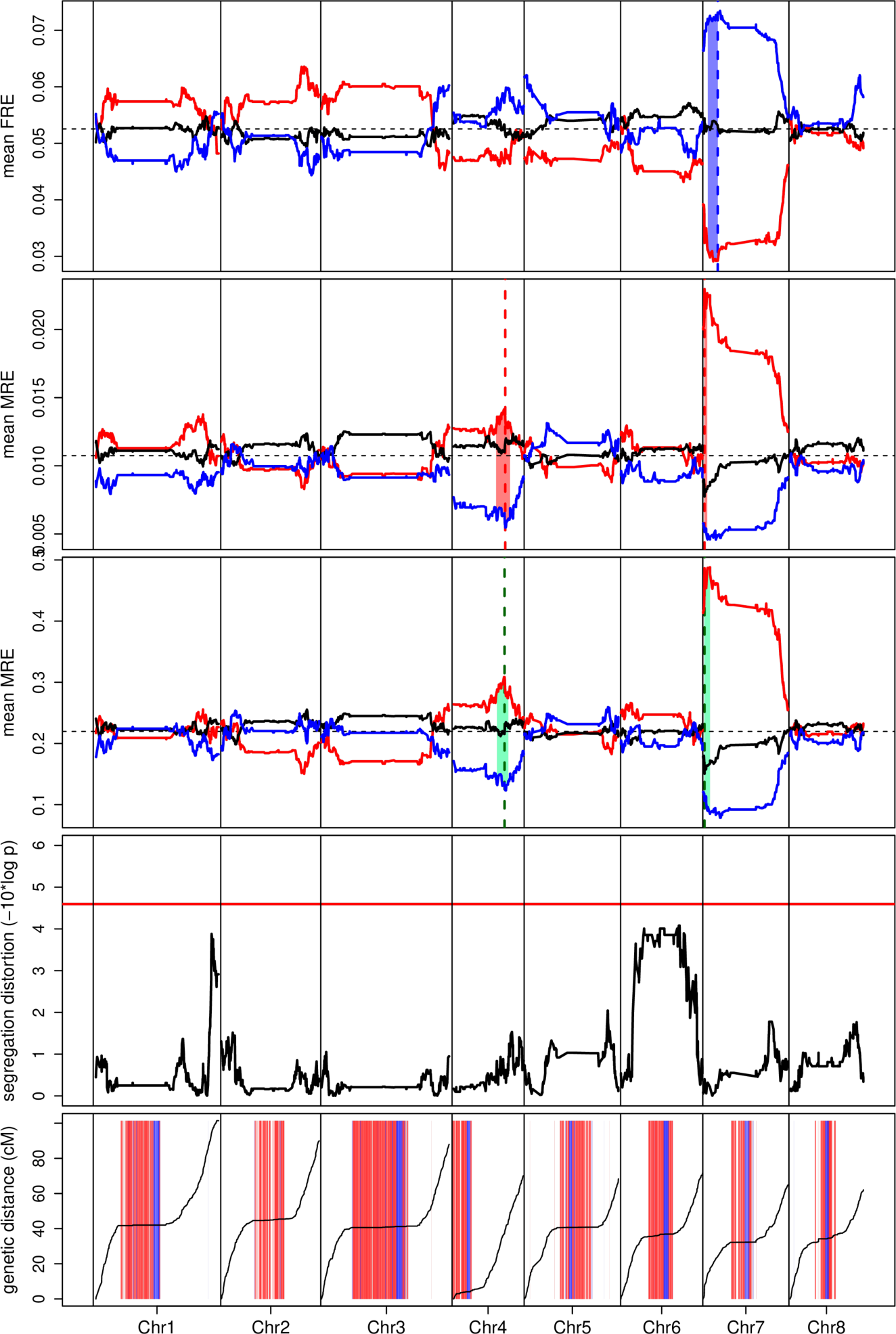
Genomic results for Cross 2. Top three plots: Mean traits (FRE, MRE, rMA) across the genome. Thick red, blue and black lines indicate mean trait values for homozygotes inherited from either Control (blue) or X-only populations (red) or heterozygotes. The horizontal, dashed line indicates mean trait values for all individuals. Vertical dashed lines indicate the location of QTL, shaded areas indicate the Bayesian credible intervals for QTL. Plot 4: *p*-value of χ²-test for segregation distortion. The red line indicates the significance threshold (*p* < 0.05) after Bonferroni correction. Bottom plot: recombination maps inferred from RADseq data. Shaded red and blue bars in the background indicate genomic windows enriched for the two most common centromeric repeats.

**Figure S8.**
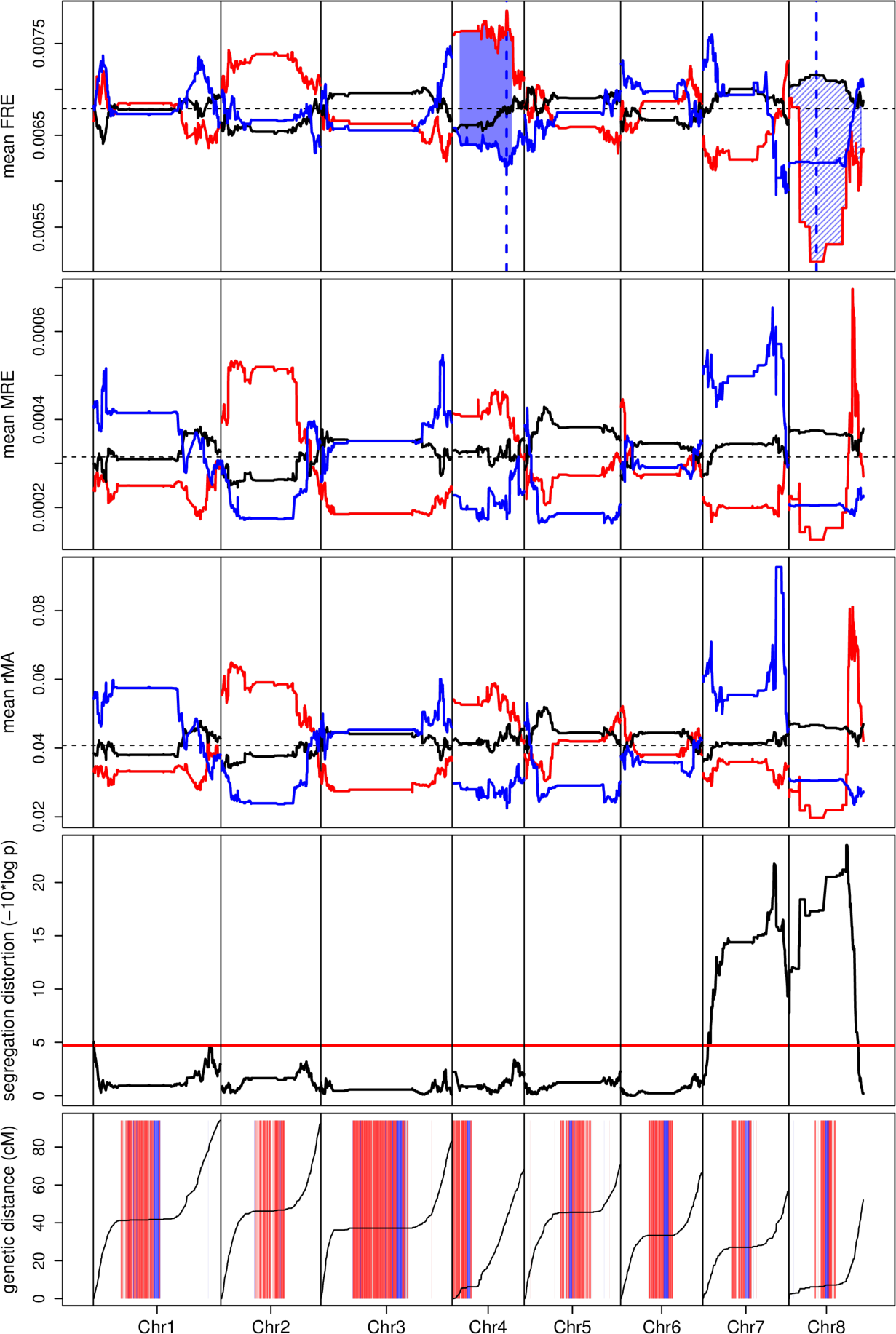
Genomic results for Cross 3. Top three plots: Mean traits (FRE, MRE, rMA) across the genome. Thick red, blue and black lines indicate mean trait values for homozygotes inherited from either Control (blue) or X-only populations (red) or heterozygotes. The horizontal, dashed line indicates mean trait values for all individuals. Vertical dashed lines indicate the location of QTL, shaded areas indicate the Bayesian credible intervals for QTL. Plot 4: *p*-value of χ²-test for segregation distortion. The red line indicates the significance threshold (*p* < 0.05) after Bonferroni correction. Bottom plot: recombination maps inferred from RADseq data. Shaded red and blue bars in the background indicate genomic windows enriched for the two most common centromeric repeats.

**Figure S9.**
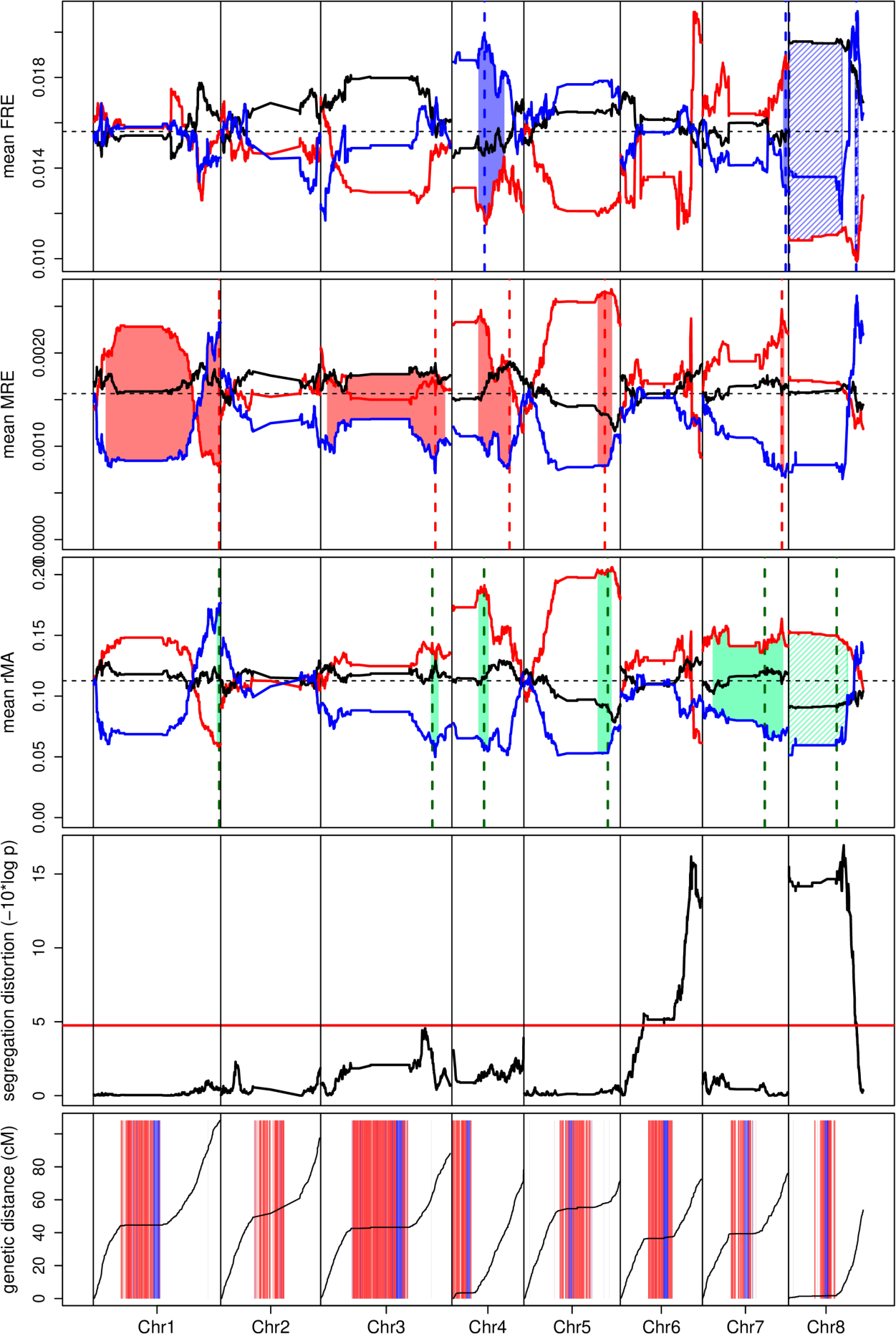
Genomic results for Cross 4. Top three plots: Mean traits (FRE, MRE, rMA) across the genome. Thick red, blue and black lines indicate mean trait values for homozygotes inherited from either Control (blue) or X-only populations (red) or heterozygotes (black). The horizontal, dashed line indicates mean trait values for all individuals. Vertical dashed lines indicate the location of QTL, shaded areas indicate the Bayesian credible intervals for QTL. Plot 4: *p*-value of χ²-test for segregation distortion. The red line indicates the significance threshold (*p* < 0.05) after Bonferroni correction. Bottom plot: recombination maps inferred from RADseq data. Shaded red and blue bars in the background indicate genomic windows enriched for the two most common centromeric repeats.

## Citations

Arends, D., P. Prins, R. C. Jansen, and K. W. Broman. 2010. R/qtl: high-throughput multiple QTL mapping. Bioinformatics 26:2990–2992.

Ashman, T. 1994. Reproductive allocation in hermaphroditic and female plants of *Sidalcea oregana* spp. *spicata* (Malvaceae) using four currencies. American Journal of Botany 81:433–438.

Ashman, T. L. 2003. Constraints on the evolution of males and sexual dimorphism: Field estimates of genetic architecture of reproductive traits in three populations of gynodioecious Fragaria virginiana. Evolution 57:2012–2025.

Ashman, T. L., and C. J. Majetic. 2006. Genetic constraints on floral evolution: a review and evaluation of patterns. Heredity 96:343–352.

Barton, N. H., and M. Turelli. 1989. Evolutionary quantitative genetics: how little do we know? Annual Review of Genetics 23:337–370.

Bawa, K. S. 1982. Seed dispersal and the evolution of dioecism in flowering plants -a response. Evolution 36:1322–1325.

Campbell, D. R. 2000. Experimental tests of sex-allocation theory in plants. Trends in Ecology & Evolution 15:227–232.

Carvalho, A. B., M. C. Sampaio, F. R. Varandas, and L. B. Klaczko. 1998. An experimental demonstration of Fisher’s principle: evolution of sexual proportion by natural selection. Genetics 148:719–731.

Catchen, J., P. A. Hohenlohe, S. Bassham, A. Amores, and W. A. Cresko. 2013. Stacks: an analysis tool set for population genomics. Molecular Ecology 22:3124–3140.

Charlesworth, D. 1999. Theories of the evolution of dioecy. Pages 33-60 in M. A. Geber, T. E. Dawson, and L. F. Delph, editors. Gender and Sexual Dimorphism in Flowering Plants. Springer, Heidelberg.

Charlesworth, D., and B. Charlesworth. 1978. A model for the evolution of dioecy and gynodioecy. American Naturalist 112:975–997.

Charlesworth, D., and B. Charlesworth. 1981. Allocation of resources to male and female functions in hermaphrodites. Biological Journal of the Linnean Society 15:57–74.

Charlesworth, D., and M. T. Morgan. 1991. Allocation of resources to sex functions in flowering plants. Philosophical Transactions of the Royal Society of London, B 332:91–102.

Charnov, E. L. 1982. The Theory of Sex Allocation. Princeton University Press, Princeton, NJ.

Charnov, E. L., J. Maynard Smith, and J. J. Bull. 1976. Why be an hermaphrodite? Nature 263:125-126.

Christenhusz, M. J. M., J. R. Pannell, A. D. Twyford, R. B. G. K. G. A. Lab, P. G. S. collective, D. T. o. L. B. collective, W. S. I. T. o. L. programme, W. S. I. S. O. D. P. collective, T. o. L. C. I. collective, and D. T. o. L. Consortiuma. 2024. The genome sequence of the annual mercury, Mercurialis annua L., 1753 (Euphorbiaceae). Wellcome Open Research 9:102.

Cossard, G., and J. R. Pannell. 2019. A functional decomposition of sex inconstancy in the dioecious, colonizing plant *Mercurialis annua*. American Journal of Botany 106:722–732.

Cossard, G. G., J. F. Gerchen, X. J. Li, Y. Cuenot, and J. R. Pannell. 2021. The rapid dissolution of dioecy by experimental evolution. Current Biology 31:1277–1283.

Cossard, G. G., and J. R. Pannell. 2021. Enhanced leaky sex expression in response to pollen limitation in the dioecious plant *Mercurialis annua*. Journal of Evolutionary Biology 34:416–422.

Cruden, R. W., and D. L. Lyon. 1985. Patterns of biomass allocation to male and female functions in plants with different mating systems. Oecologia 79:332–343.

Danecek, P., A. Auton, G. Abecasis, C. A. Albers, E. Banks, M. A. DePristo, R. E. Handsaker, G. Lunter, G. T. Marth, S. T. Sherry, G. A. T. McVean, R. Durbin, and G. P. A. Group. 2011. The variant call format and VCFtools. Bioinformatics 27:2156–2158.

Danecek, P., J. K. Bonfield, J. Liddle, J. Marshall, V. Ohan, M. O. Pollard, A. Whitwham, T. Keane, S. A. McCarthy, R. M. Davies, and H. Li. 2021. Twelve years of SAMtools and BCFtools. GigaScience 10:4.

Devisser, J., A. Termaat, and C. Zonneveld. 1994. Energy budgets and reproductive allocation in the simultaneous hermaphrodite pond snail, *Lymnaea stagnalis* (L): a trade-off between male and female function. American Naturalist 144:861–867.

Dorken, M. E., R. P. Freckleton, and J. R. Pannell. 2017. Small-scale and regional spatial dynamics of an annual plant with contrasting sexual systems. Journal of Ecology 105:1044–1057.

Eckhart, V. M. 1993. Do hermaphrodites of gynodioecious *Phacelia linearis* (Hydrophyllaceae) trade-off seed production to attract pollinators? Biological Journal of the Linnean Society 50:47–63.

Ehlers, B. K., and T. Bataillon. 2007. ’Inconstant males’ and the maintenance of labile sex expression in subdioecious plants. New Phytologist 174:194–211.

Eppley, S. M., and J. R. Pannell. 2007. Sexual systems and measures of occupancy and abundance in an annual plant: testing the metapopulation model. American Naturalist 169:20–28.

Fenster, C. B., and D. E. Carr. 1997. Genetics of sex allocation in *Mimulus* (Scrophulariaceae). Journal of Evolutionary Biology 10:641–661.

Fisher, R. A. 1930. The Genetical Theory of Natural Selection. Oxford University Press, Oxford.

Frank, S. A. 1986. Hierarchical selection theory and sex ratios 1. General solutions for structured populations. Theoretical Population Biology 29:312–342.

Frank, S. A. 2002. A touchstone in the study of adaptation. Evolution 56:2561–2564.

Friedman, J., A. J. Bohonak, and R. A. Levine. 2013. When are two pieces better than one: fitting and testing OLS and RMA regressions. Environmetrics 24:306-316.

Goldman, D. A., and M. F. Willson. 1986. Sex allocation in functionally hermaphroditic plants: A review and critique. Botanical Review 2:157–194.

Haley, C. S., and S. A. Knott. 1992. A simple regression method for mapping quantitative trait loci in line crosses using flanking markers. Heredity 69:315–324.

Hardy, I. C. W., editor. 2002. Sex Ratios: Concepts and Research Methods. Cambridge University Press, Cambridge.

Harris, M. S., and J. R. Pannell. 2008. Roots, shoots and reproduction: Sexual dimorphism in size and costs of reproductive allocation in an annual herb. Proceedings of the Royal Society of London B 275:2595–2602.

Herrera, C. M. 1982. Breeding systems and dispersal-related maternal reproductive effort of southern Spanish bird-dispersed plants. Evolution 36:1299–1314.

Hoch, J. M., and J. S. Levinton. 2012. Experimenntal tests of sex allocation theory with two species of simultaneously hermaphroditic acorn barnacles. Evolution 66:1332–1343.

Jombart, T. 2008. *adegenet*: a R package for the multivariate analysis of genetic markers. Bioinformatics 24:1403–1405.

Knaus, B. J., and N. J. Grünwald. 2017. VCFR: a package to manipulate and visualize variant call format data in R. Molecular Ecology Resources 17:44–53.

Koelewijn, H. P. 2003. Variation in restorer genes and primary sexual investment in gynodioecious Plantago coronopus: the trade-off between male and female function. Proceedings of the Royal Society of London B 270:1939–1945.

Lesaffre, T., J. R. Pannell, and C. Mullon. 2024a. An explanation for the prevalence of XY over ZW sex determination in species derived from hermaphroditism. Proceedings of the National Acadamy of Sciences USA 121:e2406305121.

Lesaffre, T., J. R. Pannell, and C. Mullon. 2024b. The joint evolution of separate sexes and sexual dimorphism. Journal of Evolutionary Biology 10.1093/jeb/voae136.

Li, H. 2013. Aligning sequence reads, clone sequences and assembly contigs with BWA-MEM. arXiv 1303.3997.

Li, X. J., P. Veltsos, G. G. Cossard, J. Gerchen, and J. R. Pannell. 2019. YY males of the dioecious plant Mercurialis annua are fully viable but produce largely infertile pollen. New Phytologist 224:1394–1404.

Locher, R., and B. Baur. 2000. Mating frequency and resource allocation to male and female function in the simultaneous hermaphrodite land snail *Arianta arbustorum*. Journal of Evolutionary Biology 13:607–614.

Mazer, S. J., V. A. Delesalle, and H. Paz. 2007. Evolution of mating system and the genetic covariance between male and female investment in *Clarkia* (Onagraceae):: Selfing opposes the evolution of trade-offs. Evolution 61:83–98.

Mölder, F., K. P. Jablonski, B. Letcher, M. B. Hall, C. H. Tomkins-Tinch, V. Sochat, J. Forster, S. Lee, S. O. Twardziok, A. Kanitz, A. Wilm, M. Holtgrewe, S. Rahmann, S. Nahnsen, and J. Köster. 2021. Sustainable data analysis with Snakemake. F1000Research 10:33.

Morgan, M. T. 1992. The evolution of traits influencing male and female fertility in outcrossing plants. American Naturalist 139:1022–1051.

Obbard, D. J., S. A. Harris, and J. R. Pannell. 2006. Sexual systems and population genetic structure in an annual plant: testing the metapopulation model. American Naturalist 167:354–366.

Pannebakker, B. A., N. Cook, J. van den Heuvel, L. van de Zande, and D. M. Shuker. 2020. Genomics of sex allocation in the parasitoid wasp *Nasonia vitripennis*. Bmc Genomics 21:14.

Pannebakker, B. A., R. Watt, S. A. Knott, S. A. West, and D. M. Shuker. 2011. The quantitative genetic basis of sex ratio variation in *Nasonia vitripennis*: a QTL study. Journal of Evolutionary Biology 24:12–22.

Pannell, J. 1997. The maintenance of gynodioecy and androdioecy in a metapopulation. Evolution 51:10–20.

Pannell, J. R. 2015. Evolution of the mating system in colonizing plants. Molecular Ecology 24:2018–2037.

Pannell, J. R., and S. C. H. Barrett. 1998. Baker’s Law revisited: reproductive assurance in a metapopulation. Evolution 52:657–668.

Paradis, E. 2010. pegas: an R package for population genetics with an integrated-modular approach. Bioinformatics 26:419–420.

Petersen, C. W. 1990. Variation in reproductive success and gonada lallocation in the simultaneous hermaphrodite, *Serranus fasciatus*. Oecologia 83:62–67.

Petersen, C. W., and E. A. Fischer. 1996. Intraspecific variation in sex allocation in a simultaneous hermaphrodite: The effect of individual size. Evolution 50:636–645.

Peterson, B. K., J. N. Weber, E. H. Kay, H. S. Fisher, and H. E. Hoekstra. 2012. Double digest RADseq: an inexpensive method for *de novo* SNP discovery and genotyping in model and non-model species. PLoS One 7:11.

Price, A. H. 2006. Believe it or not, QTLs are accurate! Trends in Plant Science 11:213–216.

Raimondi, P. T., and J. E. Martin. 1991. Evidence that mating group-size affects allocation of reproductive resources in a simultaneous hermaphrodite. American Naturalist 138:1206–1217.

Rameau, C., and P.-H. Gouyon. 1991. Resource allocation to growth, reproduction and survival in *Gladiolus*: the cost of male function. Journal of Evolutionary Biology 4:291–307.

Ramm, S. A., B. Lengerer, R. Arbore, R. Pjeta, J. Wunderer, A. Giannakara, E. Berezikov, P. Ladurner, and L. Schärer. 2019. Sex allocation plasticity on a transcriptome scale: Socially sensitive gene expression in a simultaneous hermaphrodite. Molecular Ecology 28:2321–2341.

Rastas, P. 2017. Lep-MAP3: robust linkage mapping even for low-coverage whole genome sequencing data. Bioinformatics 33:3726–3732.

Roff, D. A., and D. J. Fairbairn. 2007. The evolution of trade-offs: where are we? Journal of Evolutionary Biology 20:433–447.

Russell, J. R. W., and J. R. Pannell. 2015. Sex determination in dioecious *Mercurialis annua* and its close diploid and polyploid relatives. Heredity 114:262–271.

Schärer, L. 2009. Tests of sex allocation theory in simultaneously hermaphroditic animals. Evolution 63:1377–1405.

Schärer, L., L. M. Karlsson, M. Christen, and C. Wedekind. 2001. Size dependent sex allocation in a simultaneous hermaphrodite parasite. Journal of Evolutionary Biology 14:55–67.

Scharer, L., and P. Ladurner. 2003. Phenotypically plastic adjustment of sex allocation in a simultaneous hermaphrodite. Proceedings of the Royal Society of London B 270:935–941.

Schärer, L., P. Sandner, and N. K. Michiels. 2005. Trade-off between male and female allocation in the simultaneously hermaphroditic flatworm *Macrostomum* sp. Journal of Evolutionary Biology 18:396–404.

Seger, J., and V. M. Eckhart. 1996. Evolution of sexual systems and sex allocation in plants when growth and reproduction overlap. Proceedings of the Royal Society of London, B 263:833–841.

Smith, R. J. 2009. Use and misuse of the reduced major axis for line-fitting. American Journal of Physical Anthropology 140:476–486.

Song, S. L., and J. Z. Zhang. 2024. In search of the genetic variants of human sex ratio at birth: was Fisher wrong about sex ratio evolution? Proceedings of the Royal Society B 291:11.

Spigler, R. B., K. S. Lewers, and T. L. Ashman. 2011. Genetic architecture of sexual dimorphism in a subdioecious plant with a proto-sex chromosome. Evolution 65:1114–1126.

Toro, M. A., and B. Charlesworth. 1982. An attempt to detect genetic variation in sex ratio in *Drosophila melanogaster*. Heredity 49:199–209.

Trouvé, S., J. Jourdane, F. Renaud, P. Durand, and S. Morand. 1999. Adaptive sex allocation in a simultaneous hermaphrodite. Evolution 53:1599–1604.

van Noordwijk, A. J., and G. de Jong. 1986. Acquisition and allocation of resources: their influence on variation in life history tactics. American Naturalist 128:137–142.

Vandeputte, M., M. Dupont-Nivet, O. Merdy, P. Haffray, H. Chavanne, and B. Chatain. 2007. Quantitative genetic determinism of sex-ratio in the European sea bass (*Dicentrarchus labrax* L.). Aquaculture 272:S315–S315.

Varga, S., and M. M. Kytöviita. 2017. Lack of trade-offs between the male and female sexual functions in the gynodioecious herb *Geranium sylvaticum*. Plant Ecology 218:1163–1170.

Veltsos, P., G. Cossard, E. Beaudoing, G. Beydon, D. S. Bianchi, C. Roux, S. C. Gonzalez-Martinez, and J. R. Pannell. 2018. Size and content of the sex-determining region of the Y chromosome in dioecious *Mercurialis annua*, a plant with homomorphic sex chromosomes. Genes 9.

Veltsos, P., K. Ridout, M. A. Toups, S. C. Gonzalez-Martinez, A. Muyle, O. Emery, P. Rastas, V. Hudzieczek, R. Hobza, B. Vyskot, G. Marais, D. Filatov, and J. R. Pannell. 2019. Early sex-chromosome evolution in the diploid dioecious plant *Mercurialis annua*. Genetics 212:815–835.

West, S. A. 2009. Sex Allocation. Princeton University Press, Princeton.

West, S. A., E. A. Herre, and B. C. Sheldon. 2000. The benefits of allocating sex. Science 290:288–290.

Wlodzimierz, P., M. C. Hong, and I. R. Henderson. 2023. TRASH: tandem repeat annotation and structural hierarchy. Bioinformatics 39:8.

Zhang, D. Y., and X. H. Jiang. 2002. Size-dependent resource allocation and sex allocation in herbaceous perennial plants. Journal of Evolutionary Biology 15:74–83.

